# Resting state functional connectivity of the marmoset claustrum

**DOI:** 10.1101/2025.04.25.650655

**Authors:** Erin J. Holzscherer, Alessandro Zanini, Chun Yin Liu, Stefan Everling, David A. Seminowicz

## Abstract

The common marmoset *(Callithrix jacchus)* has been recently developed as a nonhuman primate model useful for studying behaviour, neurology, and higher-level cognitive processes considering their phylogenetic proximity to humans. Here, we investigated the resting state functional connectivity (RSFC) of the marmoset claustrum, a small, highly connected subcortical structure. Using an open resource of 234 functional MRI scans from awake marmosets, we found claustrum connectivity to the prefrontal cortex, posterior parietal cortex, temporal cortices, cingulate cortex, sensory cortices, limbic areas, basal ganglia, and cerebellum. We also found strong functional connectivity to regions and hubs involved in marmoset resting state networks. These findings demonstrate marmoset claustrum RSFC similar to previous human and rodent studies and validate the integration of marmosets into claustrum research.

## Introduction

The claustrum is a thin, grey matter structure present across mammalian brains and located between the insula and putamen. It is suggested to be the most connected region relative to its size (Torgerson et al., 2015), showing widespread bilateral projections to many cortical and subcortical regions (Arrigo et al., 2017; Borra et al., 2020, 2024; Chia et al., 2020; Fernández-Miranda et al., 2008; Gamberini et al., 2017, 2021; Gattass et al., 2014; Honda et al., 2024; Jackson et al., 2020; Krimmel et al., 2018; Krimmel et al., 2019a; Krimmel et al., 2019b; Lei et al., 2025; Milardi et al., 2015; Roberts et al., 2007; Tanné-Gariépy et al., 2002; Wang et al., 2017; White et al., 2017; White & Mathur, 2018) including those involved in major networks such as the default mode network (DMN), salience network (SN), frontoparietal network (FPN), and extrinsic mode network (EMN) (Chia et al., 2020; Jackson et al., 2020; Krimmel et al., 2018; Krimmel et al., 2019b; Madden et al., 2022; Mathur, 2014; Reser et al., 2017; Rodríguez-Vidal et al., 2024; Smith et al., 2019). Recently, the network instantiation in cognitive control (NICC) model proposed that the claustrum leverages input from frontal regions to initiate, amplify, and synchronize networks during transitions between task-positive and task-negative states (Krimmel et al., 2018; Krimmel et al., 2019b; Madden et al., 2022; Reser et al., 2017). This theory is supported by claustro-cortical connections with major hubs of functional networks (Madden et al., 2022), suggesting the claustrum supports these hubs to instantiate and synchronize networks during switching. The claustrum’s extensive connectivity and proposed role in network switching have also linked it to various neurological conditions associated with disrupted network activity, including epilepsy, Parkinson’s disease, Alzheimer’s disease, and chronic pain (Arrigo et al., 2017; Ayyildiz et al., 2023; Goll et al., 2015; Kurada et al., 2019; Ntamati et al., 2023; Stewart et al., 2024, Zedde et al., 2025). However, invasive or transgenic measures used to investigate the underlying mechanisms driving these relationships are limited by the translational value of current rodent models (Chong & Gămănuţ, 2024). While rodents can provide important mechanistic insight, studies have demonstrated primate-specific evolution of cortical neurons including differences in claustral gene (Watakabe, 2017) and cell-type (Lei et al., 2025) origins and expression patterns (Watakabe, 2017) as well as differences in primate cortical organization and higher cognition (Magrou et al., 2024). This suggests exploring a model that is more phylogenetically, neurologically and behaviourally similar to humans – such as non-human primates (NHPs) – can provide more extensive insight on the higher-level cognitive contributions of the claustrum.

The common marmoset *(Callithrix jacchus)* has become a popular translational model given its phylogenetic similarity to humans. Their small size and high fecundity give them an advantage over traditional Old-World primates while conserving many functional and anatomical brain network properties (Okano et al., 2012). Both cortical and subcortical subdivisions of the marmoset brain have been previously mapped (Paxinos et al., 2012; Saleem et al., 2024) and functional resting state networks and their major hubs (Belcher et al., 2016; Ghahremani et al., 2017, Hori et al., 2020) have been described. These networks include the DMN, SN, FPN, frontal pole network, orbitofrontal network (OFN), dorso-medial and ventral sensorimotor networks (dmSSM, vSSM), and visual networks. (Belcher et al., 2013; Burman et al., 2011; Dureux et al., 2024; Ghahremani et al., 2017; Hori et al., 2022; Hori, Schaeffer, Gilbert, et al., 2020b; Liu et al., 2019; Mitchell & Leopold, 2015). Interspecies comparisons between human and marmoset networks have revealed notable similarities in sensorimotor and visual networks, including face-, scene-, and body-specific areas (Ghahremani et al., 2017; Hori et al., 2021; Hori, Schaeffer, Yoshida, et al., 2020; Mantini et al., 2012; Ngo et al., 2023) and motion processing networks (Cloherty et al., 2020), while human frontal networks are considered a more distributed version of the ancestral frontal networks observed in marmosets (Ghahremani et al., 2017; Hori et al., 2021). Additionally, there are existing marmoset models of many of the aforementioned disorders that potentially involve aberrant claustrum activity including Parkinson’s and Alzheimer’s diseases (Okano et al., 2012; Rizzo et al., 2021; Yun et al., 2015). Given these neurological network similarities and their recent development as a disease model, the marmoset presents a promising opportunity to investigate the network contributions and disease implications of the claustrum.

Traditionally, fMRI studies investigating the claustrum have been limited by its small size, irregular shape, and proximity to the insula and putamen. Recent work in the rodent and human have addressed this limitation by developing the small region confound correction (SRCC) which removes confounding insula and putamen signal while preserving relevant claustrum signal (Krimmel et al., 2019; Krimmel et al, 2019b). Here, we investigated the resting state functional connectivity of the claustrum using SRCC in awake marmosets. We found connectivity of the claustrum with widespread cortical and subcortical areas including the marmoset DMN, FPN, frontal pole network, SN, cerebellar network, high order visual networks, cerebellum, hippocampus, amygdala and basal ganglia.

## Methods

### Data

Data from 31 awake marmosets were downloaded from an open access resource dataset (marmosetconnectome.org) (Schaeffer et al., 2022). Five animals were scanned using a 9.4 Tesla (T) 31 cm horizontal bore magnet (Varian/Agilent, Yarnton, UK) at the Centre for Functional and Metabolic Mapping at Western University (UWO) (TR = 1,500 ms, TE = 15 ms, flip angle = 35°, field of view = 64 ×64 mm, matrix size = 128 ×128, voxel size = 0.5 ×0.5 ×0.5 mm, slices = 42, bandwidth = 500 kHz, GRAPPA acceleration factor: 2 (anterior-posterior)) and 26 animals were scanned using a 7T 30 cm horizontal bore magnet (Bruker BioSpin Corp, Billerica, MA, USA) at the National Institutes for Health (NIH) (TR = 2,000 ms, TE = 22.2 ms, flip angle = 70.4°, field of view = 28 ×36 mm, matrix size = 56 ×72, voxel size = 0.5 ×0.5 ×0.5 mm, slices = 38, bandwidth = 134 kHz, GRAPPA acceleration factor: 2 (left-right)). Each animal included a T2-weighted structural image and between 6 and 22 functional scans (600 volumes each (UWO), 512 volumes each (NIH)) per animal. After exclusion of three monkeys (missing structural scans) and 6 runs (excessive noise evaluated through visual analyses), a total of 28 monkeys with 234 runs were included in the analysis. For additional details on data acquisition and scanning parameters see Schaeffer et al. (2022).

### Preprocessing

All preprocessing and registration steps were performed using Analysis of Functional NeuroImages (AFNI) (Cox, 1996) and FMRIB Software Library (FSL) (Smith et al., 2004), and preprocessing code was provided through the open access resource (Schaeffer et al., 2022). The pipeline included removing the first 10 volumes of each run to account for magnetization stabilization, despiking (AFNI 3dDespike), distortion correction (FSL topup), motion correction and censoring (AFNI 3dvolreg), and bandpass removal (0.01 to 0.1, AFNI 1dBport). Phase encoding correction was performed for data from NIH animals using left-right and right-left phase encoding collected equally across scans. Preprocessing included smoothing with a 1.5mm Gaussian kernel. However, claustrum time series was extracted from unsmoothed data. Functional scans were registered to structural images using a transformation matrix between the mean functional image and the T2-weighted image (FSL Flirt) and were normalized using a transformation matrix between the T2 image and the NIH Marmoset Brain Mapping Atlas (V3) (Advanced Normalization Tools) (Liu et al., 2021). For detailed preprocessing and registration steps, see Schaeffer et al. (2022).

### SRCC

Small region confound correction (SRCC) was used to address partial volume effects by removing “flanking” signal from regions neighbouring the claustrum (Krimmel et al., 2019b). The claustrum was dilated by 1 voxel (0.5mm) and 2 voxels (1mm) and the neighbouring region between these dilated claustrum masks were treated as noise and regressed out of the claustrum time series. All regions of interest (ROIs) were generated using the subcortical atlas of the marmoset (SAM) or the Paxinos atlas (Paxinos et al., 2012; Saleem et al., 2024). To determine which regions to include, the time series for each neighbouring region was correlated to the claustrum time series and compared to the time series of randomly selected spatially distant regions (Supplementary Table 1). Neighbouring regions were included in the correction if they had strong correlations to the claustrum prior to SRCC and weak correlations following SRCC when visually compared against the correlation of spatially distant regions. The correction included the putamen, and each subdivision of the insula (parainsular, dysgranular, agranular, granular, insular proisocortex) but excluded the amygdala. To determine the efficacy of the correction, seed-to-whole brain correlation maps were generated using the insula, putamen, and pre-SRCC claustrum as seeds. Correlation maps for each seed were inputted into a linear mixed effects model (AFNI 3dLME, Chen et al., 2013) and thresholded at the voxel level using Bonferroni correction for family-wise error (p<0.05, critical F = 25.987) and visually compared (Supplementary Figure 1).

### Statistical Analysis

Claustrum seeds were generated by extracting the left and right time series from each unsmoothened scan while removing confounding insula and putamen regions extracted during the SRCC (AFNI 3dDeconvolve, 3dmaskave). Seed-to-whole brain correlation analyses were then performed by correlating each corrected claustrum time series to the whole brain for each individual scan (n=234) while removing CSF as a nuisance regressor (AFNI 3dtcorr1d). White matter signal was not removed due to the risk of removing the claustrum signal. Correlation maps for each scan were then inputted into a linear mixed effects model (AFNI 3dLME, Chen et al., 2013) with no fixed effects and a random effect of subject to account for variance in the number of runs per animal. Resulting F values were thresholded at the voxel level using Bonferroni correction for family-wise error (p<0.05, critical F = 25.987). Grey matter activation was then associated with cortical (Paxinos et al., 2012) and subcortical (Saleem et al., 2024) regions using a custom-made Matlab script which extracted the number of activated voxels, the total number of voxels and the average F value for each cortical and subcortical ROI. The percent of activated voxels for each labeled region were calculated and activation that encompassed less than 10% of a given region was removed from further interpretation.

### Network Classification

Significant regions were classified into networks based on Belcher et al. (2013) and Ghahremani et al. (2017) (Supplementary Table 1). The following networks were included: DMN including retrosplenial cortex, posterior cingulate cortex (23, 31, 29, 30), premotor (6DR, 6Dc, 8C), medial parietal area PGM, and posterior parietal cortex (PE, PFG, PG, LIP, MIP) (Belcher et al., 2013). Dorso-medial and ventral sensorimotor networks (dmSSM/vmSSM) including primary somatosensory (1, 2, 3a-b), primary motor (4, 4c), secondary somatosensory (SP2V), cingulate cortex (23, 24) and temporoparietal regions (TPt, TPO) (Belcher et al., 2013; Ghahremani et al., 2017). Higher-order visual networks including primary visual areas, V2-V6, A19, A19M, FST, TE3, A6DM (Belcher et al., 2013). SN including anterior cingulate cortex (24), anterior insula, auditory cortex, PFG, TP, and thalamus (Belcher et al., 2013; Ghahremani et al., 2017). Orbitofrontal network including A13M, A13L, orbital periallocortex, and orbital proisocortex (Belcher et al., 2013). Frontal pole network including prefrontal cortex (8c, 9, A10) and A32 (Belcher et al., 2013). FPN including premotor (6DR, 8C), prefrontal (8aV, 8aD, 45, 47), medial parietal (PGM), and posterior parietal (PEC, LIP, VIP, MIP, AIP, PG) (Ghahremani et al., 2017). Cerebellar and subcortical areas were mapped and classified using the subcortical atlas of the marmoset (Saleem et al., 2024).

## Results

### Network connectivity

*DMN* (Figure 1A): Both the left and right claustrum seeds showed strong connectivity with regions associated with the marmoset default mode network including bilateral connectivity with the posterior parietal cortex (LIP, MIP, PG), premotor cortex (6Dr, 6Dc, 8C), and posterior cingulate cortex (23, 30). The left claustrum showed bilateral connectivity with area 31, and contralateral connectivity to PE while the right claustrum showed only ipsilateral connectivity with area 31 and no significant connectivity with PE. The right claustrum reported bilateral connectivity with area 29a-d while the left claustrum only showed contralateral connectivity in 29d.

**Figure 1.**
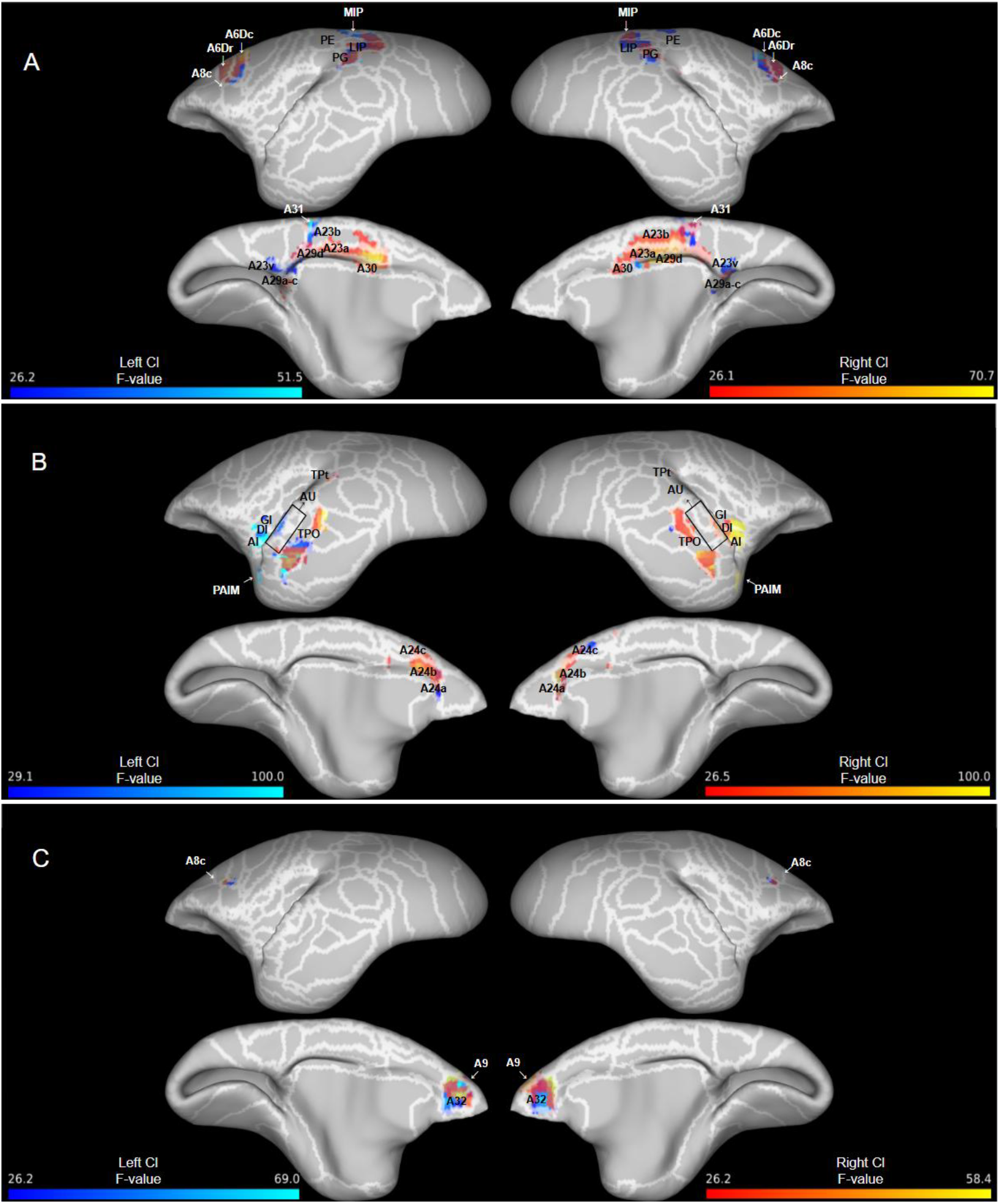
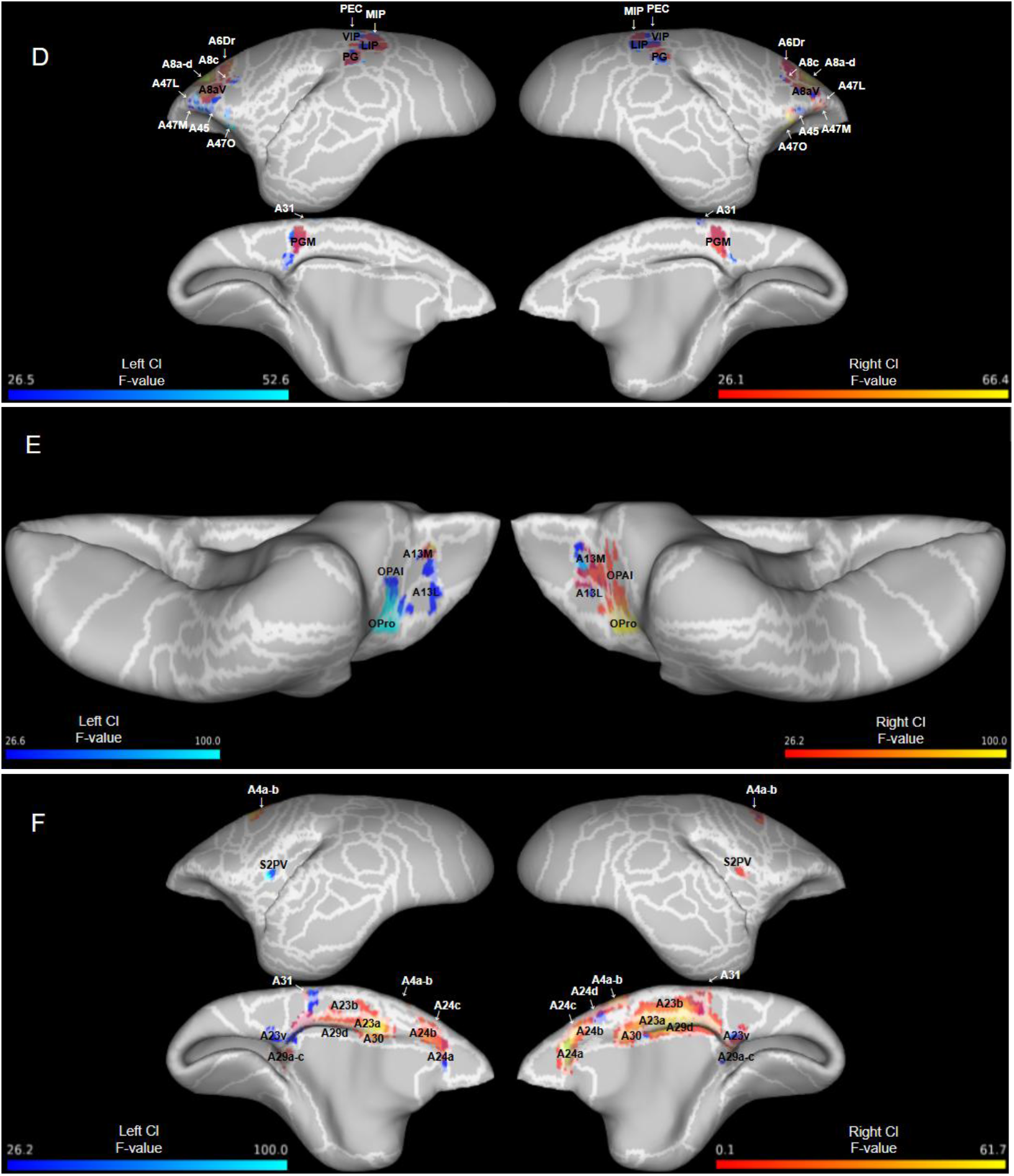
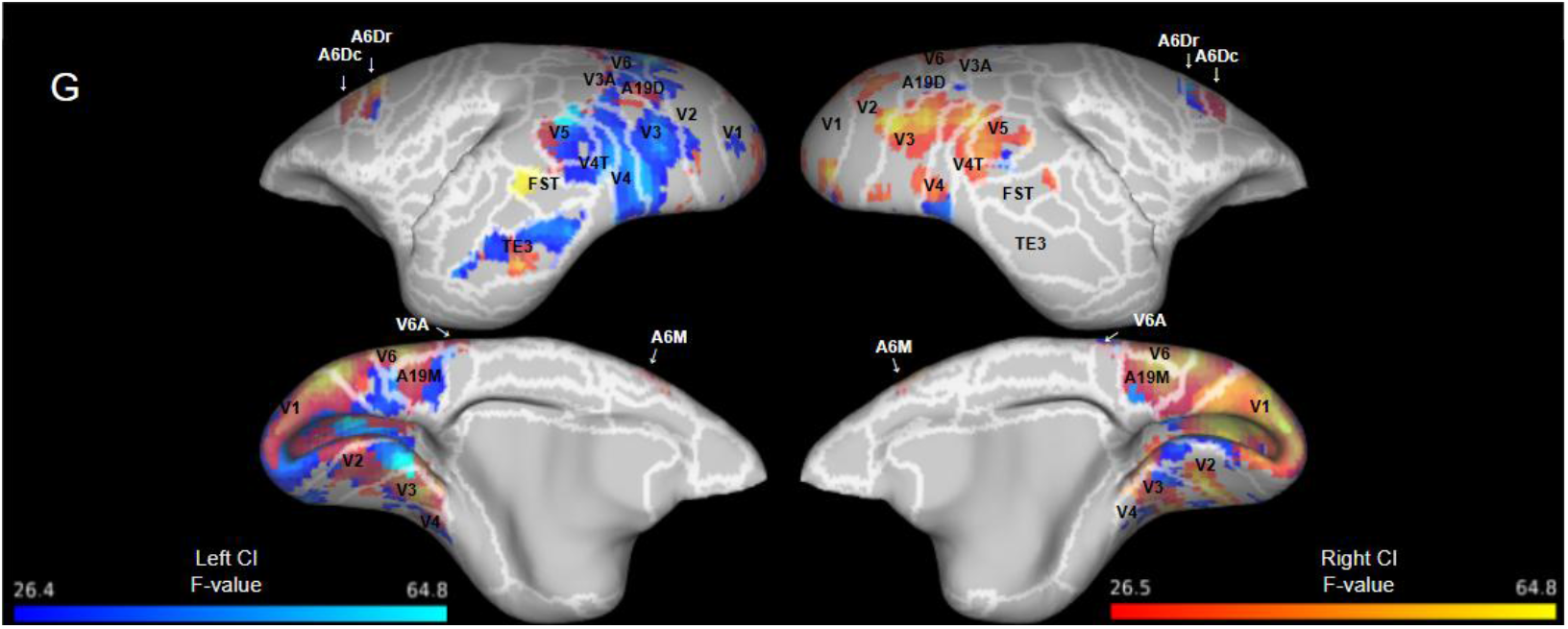
Resting state functional connectivity of both the left and right claustrum with resting state networks (RCl > LCl): (A) Default mode, (B) Salience, (C) Frontal pole, (D) Frontoparietal, (E) Orbitofrontal, (F) Sensorimotor, (G) Visual. Data presented (n=28) as significant F values (>25.987) with blue representing left claustrum connectivity and red representing the right claustrum connectivity. White lines and cortical labels delineate the Paxinos parcellation (Paxinos et al., 2012) of the NIH marmoset brain atlas (Liu et al., 2021).

*SN* (Figure 1B): Both the left and right claustrum seeds showed strong connectivity with regions associated with the marmoset salience network including bilateral connectivity with the anterior cingulate cortex (ACC, particularly in 24a-b) and ipsilateral connectivity with insular regions. The right claustrum showed bilateral connectivity with the auditory cortex, TP, and 24c while the left claustrum showed only ipsilateral connectivity with auditory cortex, and TP, and contralateral connectivity to 24d.

*Frontal Pole* (Figure 1C): Both the left and right claustrum seeds showed strong connectivity with regions associated with the marmoset frontal pole network including bilateral connectivity with the dlPFC (8c, 9) and vACC (32).

*FPN* (Figure 1D): Both the left and right claustrum seeds showed strong connectivity with regions associated with the marmoset FPN including bilateral connectivity with the posterior parietal cortex (LIP, MIP, PG, PGM), premotor cortex (6Dr, 8C), vlPFC (8aV, 8aD, 47) and ipsilateral connectivity with area 45. The left claustrum showed bilateral connectivity with PEc and VIP while the right claustrum showed only contralateral connectivity with VIP.

*OFN* (Figure 1E): Both the left and right claustrum seeds showed strong connectivity with regions associated with the marmoset orbitofrontal network including bilateral connectivity with area 13M and area 11. The left claustrum showed contralateral connectivity with area 13L while the right claustrum only showed ipsilateral connectivity. Both the left and right claustrum also showed ipsilateral connectivity to the orbital periallocortex and orbital proisocortex.

*SSM* (Figure 1F): Both the left and right claustrum seeds showed strong connectivity with regions associated with the marmoset dmSSM network but only the right claustrum showed connectivity with regions associated with the vSSM network. For the dmSSM, both left and right claustrum showed bilateral connectivity with cingulate cortex (areas 23, 24 and 30). The left claustrum showed bilateral connectivity with area 31 while the right claustrum only showed ipsilateral connectivity. Somatosensory regions (3, 1/2, S2I, S2E) involved in the dmSSM did not show significant connectivity with either claustrum seed while primary motor cortex (A4ab) showed bilateral connectivity with the right claustrum and contralateral connectivity with the left claustrum. For the vSSM, only the secondary somatosensory cortex (SP2V) showed significant ipsilateral connectivity with the left and right claustrum.

*Higher order visual* (Figure 1G): Both the left and right claustrum seeds showed strong connectivity with regions associated with the marmoset higher-order visual networks including bilateral connectivity with visual cortex (V1, V2, V3, V3A, V4, V5, V6, V6A), area 19, and area 23V. Only the right claustrum showed bilateral connectivity with FST while both the left and right claustrum showed connectivity with only the left inferior temporal cortex (TE3).

### Other cortical connectivity (Figure 2)

Both the left and right claustrum showed strong connectivity with other cortical regions that are not defined by marmoset RSNs including bilateral connectivity with dlPFC (46), mPFC (14), supplementary motor area (6M), OFC (13a-b), occipito-parietal transitional area, PGa/IPa, and vACC (25). Both the left and right claustrum showed ipsilateral connectivity with gustatory cortex, A6Va-b, secondary somatosensory cortex (S2PR), ventral temporal cortex (35, 36), and proisocortical motor region. The lateral/inferior temporal cortex showed mostly ipsilateral connectivity with both seeds while the left claustrum showed contralateral connectivity with the inferior temporal region TE1. Both claustrum seeds showed connectivity with only the left prostriate area while the entorhinal cortex and piriform cortex showed only ipsilateral connectivity with the right claustrum.

**Figure 2.**
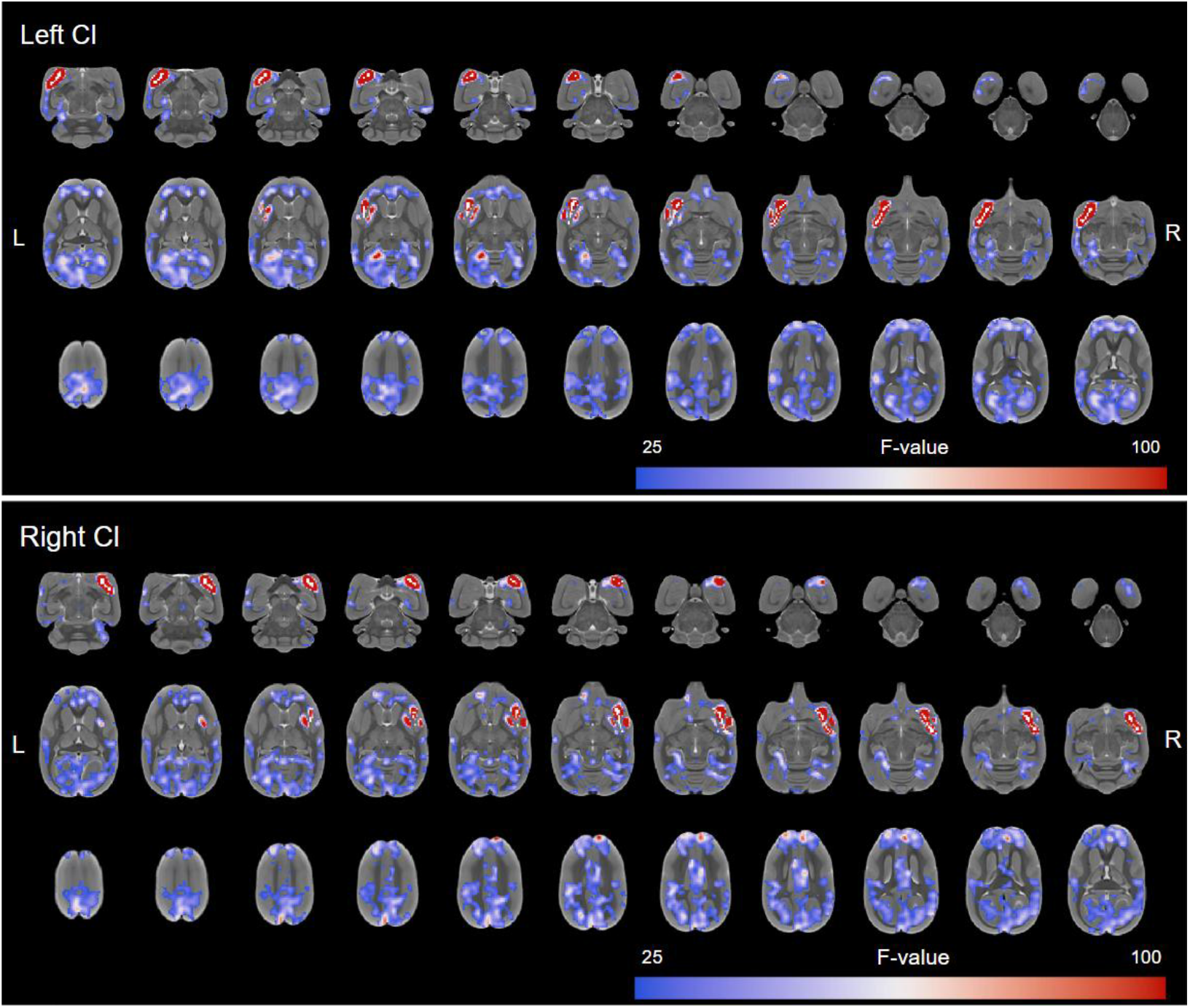
Resting state functional connectivity of the left and right claustrum with SRCC overlaid by a white claustrum mask. Data presented (n=28) as significant F values (>25.987).

### Subcortical connectivity (Figure 3)

Both the left and right claustrum showed strong ipsilateral connectivity with subcortical regions including the basal ganglia (globus pallidus, putamen), amygdala, anterior cortical nucleus, hippocampus (CA4, dentate gyrus) and olfactory tract. Both the left and right claustrum showed bilateral connectivity with the basal forebrain and subicular regions and contralateral connectivity with the superior colliculus. The right claustrum showed contralateral connectivity with the nucleus accumbens while the left claustrum showed ipsilateral connectivity with the posterior cortical nucleus and olfactory tubercle.

**Figure 3.**
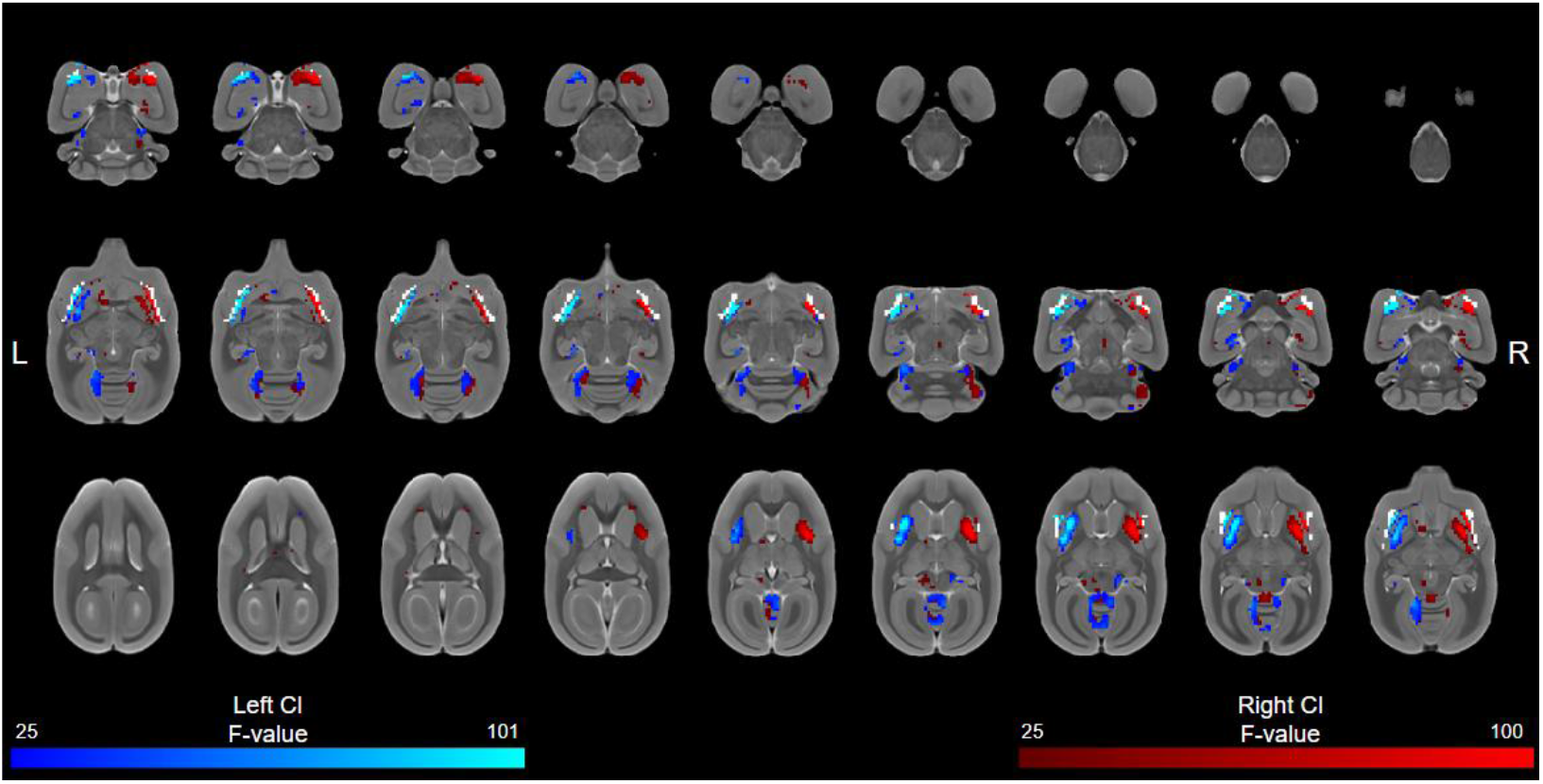
Resting state functional connectivity of both the left and right claustrum with subcortical regions (RCl>LCl) overlaid by a white claustrum mask. Data presented (n=28) as significant F values (>25.987) with blue representing left claustrum connectivity and red representing right claustrum connectivity.

Both the left and right claustrum seeds showed strong connectivity with the cerebellum including bilateral connectivity with cerebellar lobule IV and anterior part of the simple lobule. The left claustrum showed bilateral connectivity with cerebellar lobule V while the right claustrum showed only contralateral connectivity. Both seeds showed ipsilateral connectivity with the flocculus while the left claustrum showed ipsilateral connectivity with cerebellar lobule III and the right claustrum showed ipsilateral connectivity with crus I of the ansiform lobule.

For a full list of regions connected to the claustrum in the awake marmoset, see supplementary table 3.

## Discussion

The purpose of this study was to explore the resting state functional connectivity of the claustrum in the awake marmoset. Previous work has demonstrated a significant dampening effect of isoflurane on rsfMRI, especially on frontal regions of the DMN which are not activated in marmosets under anesthesia (Hori, Schaeffer, Gilbert, et al., 2020a; Smith et al., 2017). Additionally, the size and location of the claustrum has previously limited its ability to be analyzed through fMRI due to partial volume effects from neighbouring putamen and insula regions (Krimmel et al., 2019b; Madden et al., 2022; Reser et al., 2017). Here, we employed the small region confound correction, a technique used to remove confounding signals from neighbouring regions on awake marmosets to address these previous limitations (Krimmel et al., 2019b). Using 234 awake marmoset functional scans, we found left and right claustrum connectivity with prefrontal cortex, cingulate cortices, posterior parietal cortex, temporal cortex, primary sensory cortices (visual, auditory, motor), secondary sensory cortices (visual, motor, somatosensory), amygdala, basal ganglia, hippocampus and cerebellum. We found connectivity to many major regions and hubs of marmoset resting state networks including both left and right claustrum connectivity to hubs of the DMN (23b/32), salience network (24) and visual networks (V1/V2/19M/V6DM), as well as to major regions of frontal networks such as FPN (dlPFC) and orbitofrontal network (11, 13).

Generally, our findings support the claustrum’s widespread connections across many cortical and subcortical regions as seen through previous primate anatomical work. This includes regions such as prefrontal cortex, specifically A8aD and A45 (Roberts et al., 2007), anterior cingulate cortex (A24b-c, A32, A32V) (Reser et al., 2017), and hippocampus (presubiculum, parasubiculum, subiculum) (Honda et al., 2024) in the marmoset and prefrontal cortex (Lei et al., 2025; Pearson et al., 1982; Tanné-Gariépy et al., 2002), frontal-motor areas (Lei et al., 2025), parietal cortex, specifically parietal-sensory (Lei et al., 2025) and areas PEc, PGM, MIP (Gamberini et al., 2017, 2021), ventral temporal cortex (Gattass et al., 2014; Lei et al., 2025; Pearson et al., 1982), visual areas, (Gattass et al., 2014; Lei et al., 2025; Pearson et al., 1982), putamen (Borra et al., 2020; Lei et al., 2025), and amygdala (Borra et al., 2024; Lei et al., 2025) in the macaque. However, we did not find functional connectivity with thalamic nuclei which has previously shown anatomical connections in the macaque (Borra et al., 2024; Lei et al., 2025) or area 10 which has anatomical connections in the marmoset (Burman et al., 2011).

Functionally, the claustrum has been recently theorized to instantiate and synchronize functional networks as supported by connectivity to frontal and posterior hubs of both task-positive and task-negative networks (Madden et al., 2022). In line with previous work in primates and humans (Gattass et al., 2014; Jackson et al., 2020; Krimmel et al., 2018; Krimmel et al., 2019b; Lei et al., 2025; Madden et al., 2022; Pearson et al., 1982; Reser et al., 2017; Rodríguez-Vidal et al., 2024; Smith et al., 2019; Tanné-Gariépy et al., 2002), we found the marmoset claustrum showed functional connectivity with frontal and posterior hubs of marmoset specific task-positive networks (ACC, dlPFC, PPC) and task-negative networks (dlPFC, A23b, A31). Specifically, claustrum connectivity to ACC is strongly supported by anatomical (Chia et al., 2020; Reser et al., 2017; White et al., 2017; White & Mathur, 2018) and functional (Krimmel et al., 2019b; Madden et al., 2022; Rodríguez-Vidal et al., 2024) data and considered one of the main frontal hubs responsible for claustral input. Our findings are consistent with this notion considering strong bilateral connectivity between both the left and right claustrum and A24a-c, A32, A32V (Madden et al., 2022). Additionally, previous anatomical work differentiating ACC input (A24, A32) between the insula and claustrum has shown that A24d has sparse connections with the marmoset claustrum yet extensive connections with the insula (Reser et al., 2017). Here, we find functional connectivity with the claustrum and ACC is concentrated mostly in A24a-c, therefore validating the efficacy of SRCC on removing the majority of confounding insula signal. We also found strong connectivity between the marmoset claustrum and several regions in the posterior cingulate and visual cortices which are believed to receive context-dependent signals from frontal regions through the claustrum (Madden et al., 2022). The marmoset claustrum also showed significant connectivity across many regions of sensory, and frontal resting state networks, further supporting a similar role of the marmoset claustrum in cortical network regulation. These findings lay precedent for future studies to investigate the claustrum’s function and involvement in aberrant network disorders such as Parkinson’s, Alzheimer’s and chronic pain through newly developed marmoset disease models.

The marmoset claustrum also showed significant connectivity with subcortical network regions such as the amygdala, hippocampus, cerebellum and putamen. These findings are consistent with previous primate claustrum anatomical connectivity found across the hippocampus, amygdala and striatum (Borra et al., 2020, Borra et al., 2024; Gattass et al., 2014; Honda et al., 2024; Lei et al., 2025) as well as human claustro-connectivity with the hippocampus, amygdala, and basal ganglia (Fernández-Miranda et al., 2008; Krimmel et al., 2019b; Milardi et al., 2015; Rodríguez-Vidal et al., 2024). Interestingly, consistent with human findings, we found functional connectivity between the marmoset claustrum and the cerebellum (Rodríguez-Vidal et al., 2024), which is absent in rodents. In humans, this cerebellar connection has a potential role in socializing (Rodríguez-Vidal et al., 2024). In this study, this functional connectivity of the marmoset claustrum with the cerebellum, as well as with other regions critical for marmoset social processing (8b, 9) (Hori et al., 2021), may support the involvement of the claustrum in socializing processes even in this New-World non-human primate. Additionally, claustrum connectivity to the presubiculum, parasubiculum and entorhinal cortex is present both anatomically (Honda et al., 2024) and functionally in marmosets but absent in rodents. This connection is suggested to integrate memory formation with higher level cognition such as attention or self-awareness (Honda et al., 2024). These findings highlight the advantages of using the marmoset for claustrum research as their similarities in social systems, higher cognition and neural underpinnings (Miller et al., 2016) may allow for greater insight into the claustrum’s role in social network processing and memory integration.

## Limitations

While SRCC is an effective method of removing confounding insula and putamen signals, it is a relatively conservative approach and may remove relevant claustrum signals (Krimmel et al., 2019b). This possible loss of claustrum signal may explain the lack connectivity with the thalamus, hypothalamus and area 10 seen in past research in marmosets, macaques and humans (Burman et al., 2011; Honda et al., 2024; Lei et al., 2025; Madden et al., 2022) given the selective nature of claustrum projections (Lei et al., 2025) and considering connectivity to thalamic nuclei disappeared following the regression of the putamen. Future studies with increased resolution (e.g., 0.2mm isotropic) can address this limitation by reducing the amount of signal included in the regression. Additionally, because of the small, irregular size of the claustrum, data included in the time series were not smoothed and white matter was not removed during correlations, which may have introduced excess noise during seed generation and analyses. However, considering the large sample size of the data set and the consistency of our findings with previous literature, these methods likely did not have a notable effect on the cerebral connectivity pattern of the marmoset claustrum.

## Conclusion

Overall, our data demonstrates widespread claustrum connectivity with cortical and subcortical regions consistent with previous claustrum research. Given the connectivity across frontal and posterior hubs of major resting state networks, our data also suggests similar theories of claustrum function including network instantiation and task switching. However, considering marmosets and humans brains are not exact homologues and have inherent differences in cortical structure and function, caution should be used when interpreting the functional implications of these results (Kaas, 2020; Ngo et al., 2023). Future studies should investigate direct interspecies comparisons of functional connectivity using task-based methods such as movie-driven fMRI (Hori et al., 2021). Taken together, our findings act as a proof of principle by demonstrating that claustrum connectivity can be explored through a novel, phylogenetically similar animal model to facilitate future functional or disease explorations of the claustrum.

## Data and Code Availability

All raw data, preprocessing and registration code are described in Schaeffer et al. (2022) and available through the Marmoset Functional Brain Connectivity Resource (https://www.marmosetbrainconnectome.org/). SRCC and custom-made codes are available for download (https://osf.io/mga2v/?view_only=7aece5454cec49b2ae4956ad74959d4d).

## Acknowledgements

DAS was supported in part by the Wolfe-Western Fellowship at-large for outstanding newly recruited research scholars endowment fund.

## Author Contributions

**E.J.H**.: Conceptualization, Methodology, Software, Formal analysis, Writing – original draft. **A.Z**.: Methodology, Software, Formal analysis, Writing – review and editing. **C.Y.L:** Formal analysis, Writing – review and editing. **S.E**.: Resources, Writing – review and editing. **D.A.S**.: Conceptualization, Resources, Supervision, Writing – review and editing.

## References

Arrigo, A., Mormina, E., Calamuneri, A., Gaeta, M., Granata, F., Marino, S., Anastasi, G. P., Milardi, D., & Quartarone, A. (2017). Inter-hemispheric Claustral Connections in Human Brain: A Constrained Spherical Deconvolution-Based Study. Clinical Neuroradiology, 27(3), 275–281. 10.1007/s00062-015-0492-x

Ayyildiz, S., Velioglu, H. A., Ayyildiz, B., Sutcubasi, B., Hanoglu, L., Bayraktaroglu, Z., Yildirim, S., Atasever, A., & Yulug, B. (2023). Differentiation of claustrum resting-state functional connectivity in healthy aging, Alzheimer’s disease, and Parkinson’s disease. Human Brain Mapping, 44(4), 1741–1750. 10.1002/hbm.26171

Borra, E., Ballestrazzi, G., Biancheri, D., Caminiti, R., & Luppino, G. (2024). Involvement of the claustrum in the cortico-basal ganglia circuitry: connectional study in the non-human primate. Brain Structure and Function, 229(5), 1143–1164. 10.1007/s00429-024-02784-6

Borra, E., Luppino, G., Gerbella, M., Rozzi, S., & Rockland, K. S. (2020). Projections to the putamen from neurons located in the white matter and the claustrum in the macaque.

Belcher, A. M., Yen, C. C. C., Notardonato, L., Ross, T. J., Volkow, N. D., Yang, Y., Stein, E. A., Silva, A. C., & Tomasi, D. (2016). Functional connectivity hubs and networks in the awake marmoset brain. Frontiers in Integrative Neuroscience, 10(MAR2016). 10.3389/fnint.2016.00009

Belcher, A. M., Yen, C. C., Stepp, H., Gu, H., Lu, H., Yang, Y., Silva, A. C., & Stein, E. A. (2013). Large-scale brain networks in the awake, truly resting marmoset monkey. Journal of Neuroscience, 33(42), 16796–16804. 10.1523/JNEUROSCI.3146-13.2013

Burman, K. J., Reser, D. H., Richardson, K. E., Gaulke, H., Worthy, K. H., & Rosa, M. G. P. (2011). Subcortical projections to the frontal pole in the marmoset monkey. European Journal of Neuroscience, 34(2), 303–319. 10.1111/j.1460-9568.2011.07744.x

Chen, G., Saad, Z.S., Britton, J.C., Pine, D.S., Cox, R.W. (2013). Linear Mixed-Effects Modeling Approach to FMRI Group Analysis. NeuroImage 73:176–190. 10.1016/j.neuroimage.2013.01.047

Chia, Z., Augustine, G. J., & Silberberg, G. (2020). Synaptic Connectivity between the Cortex and Claustrum Is Organized into Functional Modules. Current Biology, 30(14), 2777-2790.e4. 10.1016/j.cub.2020.05.031

Chong, M. H. Y., & Gămănuţ, R. (2024). Anatomical and physiological characteristics of claustrum neurons in primates and rodents. Frontiers in Mammal Science, 3. 10.3389/fmamm.2024.1309665

Cloherty, S. L., Yates, J. L., Graf, D., Deangelis, G. C., & Mitchell, J. F. (2020). Motion Perception in the Common Marmoset. Cerebral Cortex, 30(4), 2658–2672. 10.1093/cercor/bhz267

Dureux, A., Zanini, A., Schaeffer, D. J., Johnston, K., Gilbert, K. M., & Everling, S. (2023). The marmoset default-mode network identified by deactivations in task-based fMRI studies. 10.1101/2023.08.28.555132

Fernández-Miranda, J. C., Rhoton, A. L., Kakizawa, Y., Choi, C., & Álvarez-Linera, J. (2008). The claustrum and its projection system in the human brain: A microsurgical and tractographic anatomical study - Laboratory investigation. Journal of Neurosurgery, 108(4), 764–774. 10.3171/JNS/2008/108/4/0764

Gamberini, M., Passarelli, L., Bakola, S., Impieri, D., Fattori, P., Rosa, M. G. P., & Galletti, C. (2017). Claustral afferents of superior parietal areas PEc and PE in the macaque. Journal of Comparative Neurology, 525(6), 1475–1488. 10.1002/cne.24052

Gamberini, M., Passarelli, L., Impieri, D., Montanari, G., Diomedi, S., Worthy, K. H., Burman, K. J., Reser, D. H., Fattori, P., Galletti, C., Bakola, S., & Rosa, M. G. P. (2021). Claustral Input to the Macaque Medial Posterior Parietal Cortex (Superior Parietal Lobule and Adjacent Areas). Cerebral Cortex, 31(10), 4595–4611. 10.1093/cercor/bhab108

Gattass, R., Soares, J. G. M., Desimone, R., & Ungerleider, L. G. (2014). Connectional subdivision of the claustrum: Two visuotopic subdivisions in the macaque. Frontiers in Systems Neuroscience, 8(MAY). 10.3389/fnsys.2014.00063

Ghahremani, M., Hutchison, R. M., Menon, R. S., & Everling, S. (2017). Frontoparietal functional connectivity in the common marmoset. Cerebral Cortex, 27(8), 3890–3905. 10.1093/cercor/bhw198

Goll, Y., Atlan, G., & Citri, A. (2015). Attention: The claustrum. In Trends in Neurosciences (Vol. 38, Issue 8, pp. 486–495). Elsevier Ltd. 10.1016/j.tins.2015.05.006

Hori, Y., Cléry, J. C., Schaeffer, D. J., Menon, R. S., & Everling, S. (2021). Interspecies activation correlations reveal functional correspondences between marmoset and human brain areas. 10.1101/2021.02.09.430509

Hori, Y., Clery, J. C., Schaeffer, D. J., Menon, R. S., & Everling, S. (2022). Functional Organization of Frontoparietal Cortex in the Marmoset Investigated with Awake Resting-State fMRI. Cerebral Cortex, 32(9), 1965–1977. 10.1093/cercor/bhab328

Hori, Y., Schaeffer, D. J., Gilbert, K. M., Hayrynen, L. K., Cléry, J. C., Gati, J. S., Menon, R. S., & Everling, S. (2020a). Altered Resting-State Functional Connectivity between Awake and Isoflurane Anesthetized Marmosets. Cerebral Cortex, 30(11), 5943–5959. 10.1093/cercor/bhaa168

Hori, Y., Schaeffer, D. J., Gilbert, K. M., Hayrynen, L. K., Cléry, J. C., Gati, J. S., Menon, R. S., & Everling, S. (2020b). Comparison of resting-state functional connectivity in marmosets with tracer-based cellular connectivity. NeuroImage, 204. 10.1016/j.neuroimage.2019.116241

Hori, Y., Schaeffer, D. J., Yoshida, A., Cléry, J. C., Hayrynen, L. K., Gati, J. S., Menon, R. S., & Everling, S. (2020). Cortico-subcortical functional connectivity profiles of resting-state networks in marmosets and humans. Journal of Neuroscience, 40(48), 9236–9249. 10.1523/JNEUROSCI.1984-20.2020

Jackson, J., Smith, J. B., & Lee, A. K. (2020). The Anatomy and Physiology of Claustrum-Cortex Interactions. In Annual Review of Neuroscience (Vol. 43, pp. 231–247). Annual Reviews Inc. 10.1146/annurev-neuro-092519-101637

Kaas, J. H. (2020). Comparative Functional Anatomy of Marmoset Brains. ILAR Journal, 61(2–3), 260–273. 10.1093/ilar/ilaa026

Krimmel, S. R., Qadir, H., Hesselgrave, N., White, M. G., Reser, D. H., Mathur, B. N., & Seminowicz, D. A. (2019). Resting state functional connectivity of the rat claustrum. Frontiers in Neuroanatomy, 13. 10.3389/fnana.2019.00022

Krimmel, S. R., White, M. G., Panicker, M. H., Barrett, F. S., Mathur, B. N., & Seminowicz, D. A. (2018). The human claustrum is functionally connected to cognitive networks and involved in cognitive control. 10.1101/461319

Krimmel, S. R., White, M. G., Panicker, M. H., Barrett, F. S., Mathur, B. N., & Seminowicz, D. A. (2019b). Resting state functional connectivity and cognitive task-related activation of the human claustrum. NeuroImage, 196, 59–67. 10.1016/j.neuroimage.2019.03.075

Kurada, L., Bayat, A., Joshi, S., & Koubeissi, M. Z. (2019). The claustrum in relation to seizures and electrical stimulation. In Frontiers in Neuroanatomy (Vol. 13). Frontiers Media S.A. 10.3389/fnana.2019.00008

Lei, Y., Liu, Y., Wang, M., Yuan, N., Hou, Y., Ding, L., Zhu, Z., Wu, Z., Li, C., Zheng, M., Zhang, R., Ribeiro Gomes, A. R., Xu, Y., Luo, Z., Liu, Z., Chai, Q., Misery, P., Zhong, Y., Song, X., … Shen, Z. (2025). Single-cell spatial transcriptome atlas and whole-brain connectivity of the macaque claustrum. Cell. 10.1016/j.cell.2025.02.037

Liu, C., Yen, C. C. C., Szczupak, D., Tian, X., Glen, D., & Silva, A. C. (2021). Marmoset Brain Mapping V3: Population multi-modal standard volumetric and surface-based templates. NeuroImage, 226. 10.1016/j.neuroimage.2020.117620

Liu, C., Yen, C. C. C., Szczupak, D., Ye, F. Q., Leopold, D. A., & Silva, A. C. (2019). Anatomical and functional investigation of the marmoset default mode network. Nature Communications, 10(1). 10.1038/s41467-019-09813-7

Madden, M. B., Stewart, B. W., White, M. G., Krimmel, S. R., Qadir, H., Barrett, F. S., Seminowicz, D. A., & Mathur, B. N. (2022). A role for the claustrum in cognitive control. In Trends in Cognitive Sciences (Vol. 26, Issue 12, pp. 1133–1152). Elsevier Ltd. 10.1016/j.tics.2022.09.006

Magrou, L., Joyce, M. K. P., Froudist-Walsh, S., Datta, D., Wang, X. J., Martinez-Trujillo, J., & Arnsten, A. F. T. (2024). The meso-connectomes of mouse, marmoset, and macaque: network organization and the emergence of higher cognition. Cerebral Cortex, 34(5). 10.1093/cercor/bhae174

Mantini, D., Corbetta, M., Romani, G. L., Orban, G. A., & Vanduffel, W. (2012). Data-driven analysis of analogous brain networks in monkeys and humans during natural vision. NeuroImage, 63(3), 1107–1118. 10.1016/j.neuroimage.2012.08.042

Mathur, B. N. (2014). The claustrum in review. In Frontiers in Systems Neuroscience (Vol. 8, Issue 1 APR). Frontiers Research Foundation. 10.3389/fnsys.2014.00048

Milardi, D., Bramanti, P., Milazzo, C., Finocchio, G., Arrigo, A., Santoro, G., Trimarchi, F., Quartarone, A., Anastasi, G., & Gaeta, M. (2015). Cortical and subcortical connections of the human claustrum revealed in vivo by constrained spherical deconvolution tractography. Cerebral Cortex, 25(2), 406–414. 10.1093/cercor/bht231

Miller, C. T., Freiwald, W. A., Leopold, D. A., Mitchell, J. F., Silva, A. C., & Wang, X. (2016). Marmosets: A Neuroscientific Model of Human Social Behavior. In Neuron (Vol. 90, Issue 2, pp. 219–233). Cell Press. 10.1016/j.neuron.2016.03.018

Mitchell, J. F., & Leopold, D. A. (2015). The marmoset monkey as a model for visual neuroscience. In Neuroscience Research (Vol. 93, pp. 20–46). Elsevier Ireland Ltd. 10.1016/j.neures.2015.01.008

Ngo, G. N., Hori, Y., Everling, S., & Menon, R. S. (2023). Joint-embeddings reveal functional differences in default-mode network architecture between marmosets and humans. NeuroImage, 272. 10.1016/j.neuroimage.2023.120035

Ntamati, N. R., Acuña, M. A., & Nevian, T. (2023). Pain-induced adaptations in the claustro-cingulate pathway. Cell Reports, 42(5). 10.1016/j.celrep.2023.112506

Okano, H., Hikishima, K., Iriki, A., & Sasaki, E. (2012). The common marmoset as a novel animal model system for biomedical and neuroscience research applications. In Seminars in Fetal and Neonatal Medicine (Vol. 17, Issue 6, pp. 336–340). 10.1016/j.siny.2012.07.002

Paxinos, G., Watson, C., Petrides, M., Rosa, M., Tokuno, H., 2012. The Marmoset Brain in Stereotaxic Coordinates. Elsevier, AP.

Pearson, R. C. A., Brodal, P., Gatter, K. C., & Powell, T. P. S. (1982). The organization of the connections between, the cortex and the claustrum in the monkey. In Brain Research (Vol. 234).

Reser, D. H., Majka, P., Snell, S., Chan, J. M. H., Watkins, K., Worthy, K., Quiroga, M. D. M., & Rosa, M. G. P. (2017). Topography of claustrum and insula projections to medial prefrontal and anterior cingulate cortices of the common marmoset (Callithrix jacchus). Journal of Comparative Neurology, 525(6), 1421–1441. 10.1002/cne.24009

Rizzo, S. J. S., Homanics, G. E., Park, J. E., Silva, A. C., & Strick, P. L. (2021). Establishing the marmoset as a non-human primate model of Alzheimer’s disease. Alzheimer’s & Dementia : The Journal of the Alzheimer’s Association, 17, e049952. 10.1002/alz.049952

Roberts, A. C., Tomic, D. L., Parkinson, C. H., Roeling, T. A., Cutter, D. J., Robbins, T. W., & Everitt, B. J. (2007). Forebrain connectivity of the prefrontal cortex in the marmoset monkey (Callithrix jacchus): An anterograde and retrograde tract-tracing study. Journal of Comparative Neurology, 502(1), 86–112. 10.1002/cne.21300

Rodríguez-Vidal, L., Alcauter, S., & Barrios, F. A. (2024). The functional connectivity of the human claustrum, according to the Human Connectome Project database. PLoS ONE, 19(4 April). 10.1371/journal.pone.0298349

Saleem, K. S., Avram, A. V., Glen, D., Schram, V., & Basser, P. J. (2024). The Subcortical Atlas of the Marmoset (“SAM”) monkey based on high-resolution MRI and histology. Cerebral Cortex, 34(4). 10.1093/cercor/bhae120

Schaeffer, D. J., Klassen, L. M., Hori, Y., Tian, X., Szczupak, D., Yen, C. C. C., Cléry, J. C., Gilbert, K. M., Gati, J. S., Menon, R. S., Liu, C. R., Everling, S., & Silva, A. C. (2022). An open access resource for functional brain connectivity from fully awake marmosets. NeuroImage, 252. 10.1016/j.neuroimage.2022.119030

Smith, J. B., Liang, Z., Watson, G. D. R., Alloway, K. D., & Zhang, N. (2017). Interhemispheric resting-state functional connectivity of the claustrum in the awake and anesthetized states. Brain Structure and Function, 222(5), 2041–2058. 10.1007/s00429-016-1323-9

Smith, J. B., Watson, G. D. R., Liang, Z., Liu, Y., Zhang, N., & Alloway, K. D. (2019). A role for the claustrum in salience processing? Frontiers in Neuroanatomy, 13. 10.3389/fnana.2019.00064

Stewart, B. W., Keaser, M. L., Lee, H., Margerison, S. M., Cormie, M. A., Moayedi, M., Lindquist, M. A., Chen, S., Mathur, B. N., & Seminowicz, D. A. (2024). Pathological claustrum activity drives aberrant cognitive network processing in human chronic pain. Current Biology, 34(9), 1953-1966.e6. 10.1016/j.cub.2024.03.021

Tanné-Gariépy, J., Boussaoud, D., & Rouiller, E. M. (2002). Projections of the claustrum to the primary motor, premotor, and prefrontal cortices in the macaque monkey. Journal of Comparative Neurology, 454(2), 140–157. 10.1002/cne.10425

Torgerson, C. M., Irimia, A., Goh, S. Y. M., & Van Horn, J. D. (2015). The DTI connectivity of the human claustrum. Human Brain Mapping, 36(3), 827–838. 10.1002/hbm.22667

Wang, Q., Ng, L., Harris, J. A., Feng, D., Li, Y., Royall, J. J., Oh, S. W., Bernard, A., Sunkin, S. M., Koch, C., & Zeng, H. (2017). Organization of the connections between claustrum and cortex in the mouse. Journal of Comparative Neurology, 525(6), 1317–1346. 10.1002/cne.24047

Watakabe, A. (2017). In situ hybridization analyses of claustrum-enriched genes in marmosets. Journal of Comparative Neurology, 525(6), 1442–1458. 10.1002/cne.24021

White, M. G., Cody, P. A., Bubser, M., Wang, H. D., Deutch, A. Y., & Mathur, B. N. (2017). Cortical hierarchy governs rat claustrocortical circuit organization. Journal of Comparative Neurology, 525(6), 1347–1362. 10.1002/cne.23970

White, M. G., & Mathur, B. N. (2018). Frontal cortical control of posterior sensory and association cortices through the claustrum. Brain Structure and Function, 223(6), 2999–3006. 10.1007/s00429-018-1661-x

Yun, J. W., Ahn, J. B., & Kang, B. C. (2015). Modeling Parkinson’s disease in the common marmoset (Callithrix jacchus): overview of models, methods, and animal care. In Laboratory Animal Research (Vol. 31, Issue 4, pp. 155–165). BioMed Central Ltd. 10.5625/lar.2015.31.4.155

Zedde, M., Quatrale, R., Cossu, G., Sette, M. Del, & Pascarella, R. (2025). The Role of the Claustrum in Parkinson’s Disease and Vascular Parkinsonism: A Matter of Network? In Life (Vol. 15, Issue 2). Multidisciplinary Digital Publishing Institute (MDPI). 10.3390/life15020180

