## Supplementary Material for "Resting state functional connectivity of the marmoset claustrum"

**Table 1.** Correlations between the claustrum time series and the time series of ipsilateral neighbouring regions (putamen, amygdala and insula subdivisions including: agranular, dysgranular, granular, parainsular, prosiocortex) and spatially distant regions (cortical: V2 and subcortical: ventral pallidum).

| Region | Left Time Series Correlation | Right Time Series Correlation |
| --- | --- | --- |
| Agranular | 0.555089 | 0.562642 |
| Dysgranular | 0.587848 | 0.578302 |
| Granular | 0.524242 | 0.497207 |
| Parainsular | 0.463241 | 0.464980 |
| Proisocortex | 0.602810 | 0.572881 |
| Temporal Proisocortex | 0.602810 | 0.572881 |
| Putamen | 0.501238 | 0.446953 |
| Amygdala | 0.222388 | 0.271033 |
| Ventral pallidum | 0.150012 | 0.150252 |
| V2 | 0.193217 | 0.259509 |

**Table 2.** Networks defined by Belcher et al. (2013) and Ghahremani et al. (2017) and their associated regions. Masks were created using the Paxinos atlas (Paxinos et al., 2012).

| Network | Regions | Mask |
|---|---|---|
| Default Mode | Retrosplenial cortex<br>Posterior cingulate (23, 31, 29, 30)<br>Premotor (6DR, 6Dc, 8C)<br>Medial parietal area (PGM)<br>Posterior parietal cortex (PE, PFG, PG, LIP, MIP) | 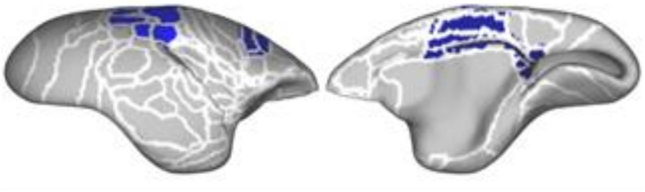 |
| Salience | Anterior cingulate (24)<br>Anterior insula<br>Auditory cortex<br>PFG<br>TP<br>Thalamus | 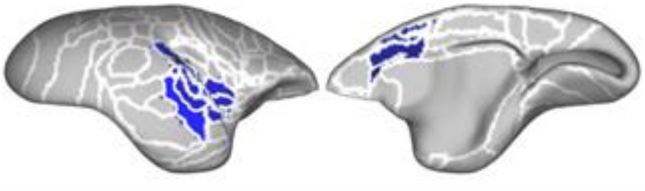 |
| Orbitofrontal | A13M<br>A13L<br>Orbital periallocortex<br>Orbital proisocortex | 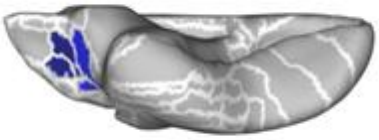 |
| Fronto-parietal | Premotor (6DR, 8C)<br>PFC (8aV, 8aD, 45, 47)<br>Medial parietal (PGM)<br>Posterior parietal (PEC, LIP, VIP, MIP, AIP, PG) | 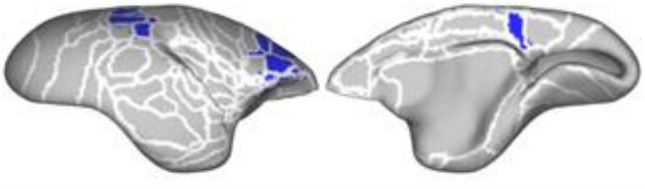 |
| Frontal pole | PFC (A8c, A9, A10)<br>A32 | 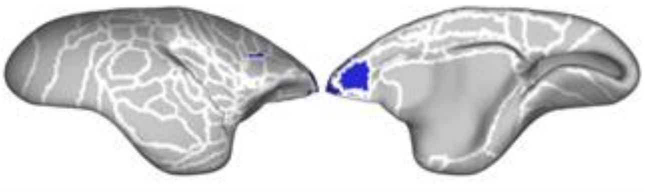 |

| | | |
|---|---|---|
| Visual | V1-V6<br>A19<br>A19M<br>FST<br>TE3<br>A6DM | 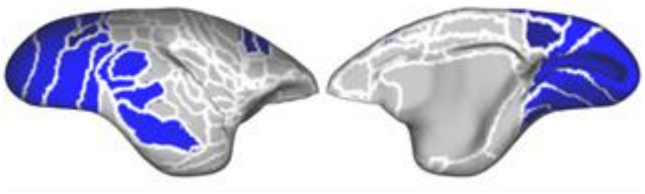 |
| Sensorimotor | S1 (1, 2, 3a-b)<br>M1 (4, 4c)<br>S2 (SP2V)<br>Cingulate cortex (23, 24)<br>Temporoparietal regions (TPt, TPO) | 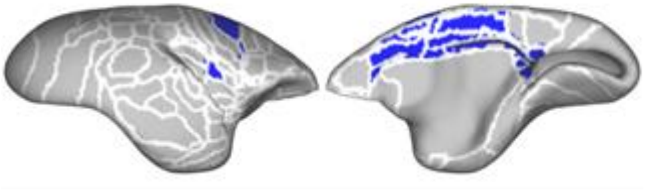 |

**Table 3.** The average F value and the percent of significant voxels for cortical (Paxinos atlas, Paxinos et al., 2012) and subcortical (subcortical atlas of the marmoset (SAM), Saleem et al., 2024) regions for each claustrum (left, right) and projection (ipsilateral, contralateral).

| Average F | Percent Activated | Claustrum | Projection | Atlas | Region |
|---|---|---|---|---|---|
| 40.296 | 28.448 | Left | Ipsilateral | SAM | globus pallidus external segment |
| 40.695 | 26.724 | Right | Ipsilateral | SAM | globus pallidus external segment |
| 34.050 | 10.112 | Right | Contralateral | SAM | nucleus accumbens |
| 50.313 | 21.429 | Left | Ipsilateral | SAM | olfactory tubercle |
| 69.077 | 53.388 | Left | Ipsilateral | SAM | Putamen |
| 69.931 | 52.772 | Right | Ipsilateral | SAM | Putamen |
| 30.178 | 16.667 | Left | Contralateral | SAM | anterior part of the simple lobule |
| 32.146 | 34.314 | Left | Ipsilateral | SAM | anterior part of the simple lobule |
| 29.087 | 12.745 | Right | Contralateral | SAM | anterior part of the simple lobule |
| 39.312 | 49.020 | Right | Ipsilateral | SAM | anterior part of the simple lobule |
| 32.525 | 11.765 | Left | Ipsilateral | SAM | cerebellar lobule III |
| 36.286 | 35.065 | Left | Contralateral | SAM | cerebellar lobule IV |
| 38.533 | 44.156 | Left | Ipsilateral | SAM | cerebellar lobule IV |
| 41.960 | 41.558 | Right | Contralateral | SAM | cerebellar lobule IV |
| 33.422 | 17.532 | Right | Ipsilateral | SAM | cerebellar lobule IV |
| 33.722 | 19.304 | Left | Contralateral | SAM | cerebellar lobule V |
| 35.980 | 32.278 | Left | Ipsilateral | SAM | cerebellar lobule V |
| 35.131 | 27.532 | Right | Contralateral | SAM | cerebellar lobule V |
| 36.242 | 39.726 | Right | Ipsilateral | SAM | crus I of the ansiform lobule |
| 30.430 | 10.497 | Left | Contralateral | SAM | flocculus |
| 36.840 | 29.282 | Left | Ipsilateral | SAM | flocculus |
| 32.682 | 14.917 | Right | Ipsilateral | SAM | flocculus |
| 35.927 | 21.374 | Right | Ipsilateral | SAM | posterior part of the simple lobule |
| 32.751 | 34.615 | Right | Ipsilateral | Paxinos | amygdalopiriform transition area |
| 35.470 | 47.748 | Left | Contralateral | Paxinos | area 11 of cortex |
| 35.392 | 23.423 | Left | Ipsilateral | Paxinos | area 11 of cortex |
| 50.697 | 67.568 | Right | Contralateral | Paxinos | area 11 of cortex |
| 34.110 | 26.126 | Right | Ipsilateral | Paxinos | area 11 of cortex |
| 38.794 | 52.381 | Left | Contralateral | Paxinos | area 13a of cortex |
| 38.173 | 19.048 | Left | Ipsilateral | Paxinos | area 13a of cortex |
| 37.175 | 47.619 | Right | Contralateral | Paxinos | area 13a of cortex |
| 29.671 | 23.810 | Right | Ipsilateral | Paxinos | area 13a of cortex |
| 42.989 | 93.103 | Left | Contralateral | Paxinos | area 13b of cortex |
| 43.578 | 17.241 | Left | Ipsilateral | Paxinos | area 13b of cortex |
| 49.498 | 68.966 | Right | Contralateral | Paxinos | area 13b of cortex |
| 30.850 | 24.138 | Right | Ipsilateral | Paxinos | area 13b of cortex |
| 28.623 | 16.667 | Left | Ipsilateral | Paxinos | area 14 of cortex caudal part |
| 32.514 | 33.333 | Right | Contralateral | Paxinos | area 14 of cortex caudal part |
| 36.806 | 17.284 | Left | Contralateral | Paxinos | area 14 of cortex rostral part |
| 33.116 | 13.580 | Right | Contralateral | Paxinos | area 14 of cortex rostral part |
| 32.440 | 20.000 | Left | Contralateral | Paxinos | area 25 of cortex |
| 35.704 | 61.429 | Left | Ipsilateral | Paxinos | area 25 of cortex |
| 30.548 | 11.429 | Right | Contralateral | Paxinos | area 25 of cortex |
| 32.408 | 25.714 | Right | Ipsilateral | Paxinos | area 25 of cortex |
| 54.834 | 27.660 | Left | Ipsilateral | Paxinos | area 35 of cortex |
| 50.941 | 36.170 | Right | Ipsilateral | Paxinos | area 35 of cortex |
| 41.892 | 39.037 | Left | Ipsilateral | Paxinos | area 36 of cortex |
| 54.568 | 17.647 | Right | Ipsilateral | Paxinos | area 36 of cortex |
| 38.182 | 50.000 | Left | Ipsilateral | Paxinos | area 46 of cortex dorsal part |
| 34.541 | 61.538 | Right | Contralateral | Paxinos | area 46 of cortex dorsal part |
| 34.720 | 57.692 | Right | Ipsilateral | Paxinos | area 46 of cortex dorsal part |
| 32.339 | 36.585 | Left | Contralateral | Paxinos | area 46 of cortex ventral part |
| 30.870 | 58.537 | Left | Ipsilateral | Paxinos | area 46 of cortex ventral part |
| 33.092 | 41.463 | Right | Contralateral | Paxinos | area 46 of cortex ventral part |
| 34.914 | 60.976 | Right | Ipsilateral | Paxinos | area 46 of cortex ventral part |
| 39.961 | 67.568 | Left | Contralateral | Paxinos | area 6 of cortex medial (supplementary motor) part |
| 36.603 | 59.459 | Left | Ipsilateral | Paxinos | area 6 of cortex medial (supplementary motor) part |
| 40.416 | 79.279 | Right | Contralateral | Paxinos | area 6 of cortex medial (supplementary motor) part |
| 38.822 | 76.577 | Right | Ipsilateral | Paxinos | area 6 of cortex medial (supplementary motor) part |
| 40.402 | 16.484 | Left | Ipsilateral | Paxinos | area 6 of cortex ventral part a |
| 53.116 | 10.989 | Right | Ipsilateral | Paxinos | area 6 of cortex ventral part a |
| 81.775 | 51.613 | Left | Ipsilateral | Paxinos | area 6 of cortex ventral part b |
| 81.619 | 45.161 | Right | Ipsilateral | Paxinos | area 6 of cortex ventral part b |
| 35.544 | 21.212 | Right | Ipsilateral | Paxinos | entorhinal cortex |
| 36.705 | 56.044 | Left | Contralateral | Paxinos | occipito-parietal transitional area of cortex |
| 40.676 | 100.000 | Left | Ipsilateral | Paxinos | occipito-parietal transitional area of cortex |
| 41.201 | 61.538 | Right | Contralateral | Paxinos | occipito-parietal transitional area of cortex |
| 28.750 | 15.385 | Right | Ipsilateral | Paxinos | occipito-parietal transitional area of cortex |
| 33.184 | 33.333 | Left | Contralateral | Paxinos | parietal areas PGa and IPa (fundus of superior temporal ventral area) |

| | | | | | |
|---|---|---|---|---|---|
| 32.004 | 25.758 | Left | Ipsilateral | Paxinos | parietal areas PGa and IPa (fundus of superior temporal ventral area) |
| 29.691 | 22.727 | Right | Contralateral | Paxinos | parietal areas PGa and IPa (fundus of superior temporal ventral area) |
| 30.007 | 33.333 | Right | Ipsilateral | Paxinos | parietal areas PGa and IPa (fundus of superior temporal ventral area) |
| 84.845 | 53.774 | Left | Ipsilateral | Paxinos | piriform cortex |
| 74.335 | 65.094 | Right | Ipsilateral | Paxinos | piriform cortex |
| 70.168 | 36.111 | Left | Ipsilateral | Paxinos | proisocortical motor region (precentral opercular cortex) |
| 80.296 | 40.741 | Right | Ipsilateral | Paxinos | proisocortical motor region (precentral opercular cortex) |
| 29.295 | 10.870 | Left | Contralateral | Paxinos | prostriate area |
| 28.309 | 19.565 | Left | Ipsilateral | Paxinos | prostriate area |
| 34.836 | 47.826 | Right | Contralateral | Paxinos | prostriate area |
| 75.919 | 33.333 | Left | Ipsilateral | Paxinos | superior temporal rostral area (cortex) |
| 72.424 | 27.193 | Right | Ipsilateral | Paxinos | superior temporal rostral area (cortex) |
| 36.409 | 27.119 | Left | Contralateral | Paxinos | temporal area TE occipital part |
| 34.708 | 36.017 | Left | Ipsilateral | Paxinos | temporal area TE occipital part |
| 57.515 | 47.287 | Left | Ipsilateral | Paxinos | temporal area TE1 (inferior temporal cortex) |
| 81.602 | 23.256 | Right | Ipsilateral | Paxinos | temporal area TE1 (inferior temporal cortex) |
| 38.796 | 60.294 | Left | Contralateral | Paxinos | temporal area TF occipital part |
| 32.164 | 20.889 | Left | Ipsilateral | Paxinos | temporal area TH |
| 32.354 | 25.926 | Left | Contralateral | Paxinos | temporal area TL occipital part |
| 38.646 | 69.466 | Left | Contralateral | Paxinos | area 6 of cortex dorsocaudal part |
| 31.768 | 46.565 | Left | Ipsilateral | Paxinos | area 6 of cortex dorsocaudal part |
| 43.966 | 60.305 | Right | Contralateral | Paxinos | area 6 of cortex dorsocaudal part |
| 32.211 | 32.824 | Right | Ipsilateral | Paxinos | area 6 of cortex dorsocaudal part |
| 30.215 | 53.788 | Left | Contralateral | Paxinos | lateral intraparietal area of cortex |
| 30.750 | 46.212 | Left | Ipsilateral | Paxinos | lateral intraparietal area of cortex |
| 30.345 | 52.273 | Right | Contralateral | Paxinos | lateral intraparietal area of cortex |
| 27.463 | 20.455 | Right | Ipsilateral | Paxinos | lateral intraparietal area of cortex |
| 29.156 | 61.111 | Left | Contralateral | Paxinos | medial intraparietal area of cortex |
| 33.947 | 84.722 | Left | Ipsilateral | Paxinos | medial intraparietal area of cortex |
| 27.944 | 47.222 | Right | Contralateral | Paxinos | medial intraparietal area of cortex |
| 29.546 | 43.056 | Right | Ipsilateral | Paxinos | medial intraparietal area of cortex |
| 28.565 | 15.110 | Left | Contralateral | Paxinos | parietal area PE |
| 33.730 | 12.879 | Left | Contralateral | SAM | superior colliculus |
| 28.700 | 21.970 | Right | Contralateral | SAM | superior colliculus |
| 32.156 | 17.763 | Left | Contralateral | Paxinos | area 23a of cortex |
| 35.702 | 21.711 | Left | Ipsilateral | Paxinos | area 23a of cortex |
| 39.613 | 76.974 | Right | Ipsilateral | Paxinos | area 23a of cortex |
| 34.339 | 33.108 | Left | Ipsilateral | Paxinos | area 23b of cortex |
| 30.156 | 47.297 | Right | Contralateral | Paxinos | area 23b of cortex |
| 30.871 | 37.162 | Right | Ipsilateral | Paxinos | area 23b of cortex |
| 32.965 | 11.765 | Right | Contralateral | Paxinos | area 23c of cortex |
| 32.752 | 42.105 | Right | Contralateral | Paxinos | area 29a-c of cortex |
| 35.585 | 20.000 | Left | Contralateral | Paxinos | area 29d of cortex |
| 33.121 | 25.455 | Right | Contralateral | Paxinos | area 29d of cortex |
| 42.651 | 54.545 | Right | Ipsilateral | Paxinos | area 29d of cortex |
| 33.545 | 14.835 | Left | Contralateral | Paxinos | area 30 of cortex |
| 29.704 | 23.626 | Left | Ipsilateral | Paxinos | area 30 of cortex |
| 34.521 | 46.154 | Right | Contralateral | Paxinos | area 30 of cortex |
| 40.738 | 60.440 | Right | Ipsilateral | Paxinos | area 30 of cortex |
| 32.974 | 35.294 | Left | Contralateral | Paxinos | area 31 of cortex |
| 39.704 | 44.706 | Left | Ipsilateral | Paxinos | area 31 of cortex |
| 28.657 | 21.176 | Right | Ipsilateral | Paxinos | area 31 of cortex |
| 33.041 | 75.789 | Left | Contralateral | Paxinos | area 6 of cortex dorsorostral part |
| 32.882 | 71.579 | Left | Ipsilateral | Paxinos | area 6 of cortex dorsorostral part |
| 39.738 | 88.421 | Right | Contralateral | Paxinos | area 6 of cortex dorsorostral part |
| 36.146 | 66.316 | Right | Ipsilateral | Paxinos | area 6 of cortex dorsorostral part |
| 29.426 | 30.851 | Left | Contralateral | Paxinos | parietal area PG |
| 33.899 | 71.277 | Left | Ipsilateral | Paxinos | parietal area PG |
| 36.095 | 44.681 | Right | Contralateral | Paxinos | parietal area PG |
| 30.481 | 38.298 | Right | Ipsilateral | Paxinos | parietal area PG |
| 32.000 | 21.053 | Left | Contralateral | Paxinos | area 8 of cortex caudal part |
| 35.674 | 50.000 | Left | Ipsilateral | Paxinos | area 8 of cortex caudal part |
| 28.773 | 26.316 | Right | Contralateral | Paxinos | area 8 of cortex caudal part |
| 35.596 | 26.316 | Right | Ipsilateral | Paxinos | area 8 of cortex caudal part |
| 32.196 | 79.032 | Left | Contralateral | Paxinos | area 23 of cortex ventral part |
| 33.452 | 67.742 | Left | Ipsilateral | Paxinos | area 23 of cortex ventral part |
| 31.980 | 37.097 | Right | Contralateral | Paxinos | area 23 of cortex ventral part |
| 29.438 | 32.258 | Right | Ipsilateral | Paxinos | area 23 of cortex ventral part |
| 37.727 | 104.605 | Right | Contralateral | Paxinos | area 23a of cortex |
| 35.800 | 10.565 | Left | Contralateral | Paxinos | area 4 of cortex parts a and b (primary motor) |
| 39.459 | 21.867 | Right | Contralateral | Paxinos | area 4 of cortex parts a and b (primary motor) |
| 31.581 | 24.570 | Right | Ipsilateral | Paxinos | area 4 of cortex parts a and b (primary motor) |
| 43.008 | 70.796 | Left | Contralateral | Paxinos | area 32 of cortex |
| 39.990 | 100.000 | Left | Ipsilateral | Paxinos | area 32 of cortex |
| 33.465 | 76.106 | Right | Contralateral | Paxinos | area 32 of cortex |

| | | | | | |
|---|---|---|---|---|---|
| 34.307 | 48.673 | Right | Ipsilateral | Paxinos | area 32 of cortex |
| 41.189 | 57.143 | Left | Contralateral | Paxinos | area 32 of cortex ventral part |
| 33.892 | 61.905 | Left | Ipsilateral | Paxinos | area 32 of cortex ventral part |
| 29.453 | 19.048 | Right | Contralateral | Paxinos | area 32 of cortex ventral part |
| 34.388 | 56.667 | Left | Contralateral | Paxinos | area 8b of cortex |
| 41.009 | 81.111 | Left | Ipsilateral | Paxinos | area 8b of cortex |
| 42.814 | 101.111 | Right | Contralateral | Paxinos | area 8b of cortex |
| 60.376 | 85.556 | Right | Ipsilateral | Paxinos | area 8b of cortex |
| 31.285 | 15.294 | Left | Contralateral | Paxinos | area 9 of cortex |
| 45.671 | 60.000 | Left | Ipsilateral | Paxinos | area 9 of cortex |
| 33.818 | 58.824 | Right | Contralateral | Paxinos | area 9 of cortex |
| 42.468 | 74.118 | Right | Ipsilateral | Paxinos | area 9 of cortex |
| 30.071 | 11.429 | Left | Contralateral | Paxinos | area 45 of cortex |
| 34.667 | 25.714 | Left | Ipsilateral | Paxinos | area 45 of cortex |
| 40.770 | 11.429 | Right | Ipsilateral | Paxinos | area 45 of cortex |
| 36.177 | 12.950 | Left | Ipsilateral | Paxinos | area 47 (old 12) of cortex lateral part |
| 39.966 | 25.899 | Right | Ipsilateral | Paxinos | area 47 (old 12) of cortex lateral part |
| 33.563 | 19.565 | Left | Contralateral | Paxinos | area 47 (old 12) of cortex medial part |
| 41.045 | 45.652 | Left | Ipsilateral | Paxinos | area 47 (old 12) of cortex medial part |
| 41.790 | 19.565 | Right | Ipsilateral | Paxinos | area 47 (old 12) of cortex medial part |
| 29.981 | 13.333 | Left | Contralateral | Paxinos | area 47 (old 12) of cortex orbital part |
| 62.263 | 42.222 | Left | Ipsilateral | Paxinos | area 47 (old 12) of cortex orbital part |
| 59.826 | 53.333 | Right | Ipsilateral | Paxinos | area 47 (old 12) of cortex orbital part |
| 35.086 | 68.345 | Left | Contralateral | Paxinos | area 8a of cortex dorsal part |
| 41.142 | 94.245 | Left | Ipsilateral | Paxinos | area 8a of cortex dorsal part |
| 47.427 | 76.978 | Right | Contralateral | Paxinos | area 8a of cortex dorsal part |
| 46.663 | 95.683 | Right | Ipsilateral | Paxinos | area 8a of cortex dorsal part |
| 42.308 | 51.449 | Left | Contralateral | Paxinos | area 8a of cortex ventral par |
| 43.124 | 69.565 | Left | Ipsilateral | Paxinos | area 8a of cortex ventral par |
| 31.077 | 52.899 | Right | Contralateral | Paxinos | area 8a of cortex ventral par |
| 35.653 | 44.928 | Right | Ipsilateral | Paxinos | area 8a of cortex ventral par |
| 28.501 | 47.458 | Left | Contralateral | Paxinos | parietal area PE caudal part |
| 39.064 | 72.881 | Left | Ipsilateral | Paxinos | parietal area PE caudal part |
| 31.896 | 49.425 | Left | Contralateral | Paxinos | parietal area PG medial part (cortex) |
| 32.869 | 120.690 | Left | Ipsilateral | Paxinos | parietal area PG medial part (cortex) |
| 29.609 | 65.517 | Right | Contralateral | Paxinos | parietal area PG medial part (cortex) |
| 31.190 | 54.023 | Right | Ipsilateral | Paxinos | parietal area PG medial part (cortex) |
| 27.368 | 16.667 | Left | Contralateral | Paxinos | ventral intraparietal area of cortex |
| 28.547 | 33.333 | Left | Ipsilateral | Paxinos | ventral intraparietal area of cortex |
| 29.618 | 46.667 | Right | Contralateral | Paxinos | ventral intraparietal area of cortex |
| 33.039 | 24.942 | Left | Contralateral | Paxinos | visual area 2 |
| 37.466 | 47.397 | Left | Ipsilateral | Paxinos | visual area 2 |
| 36.911 | 31.158 | Right | Contralateral | Paxinos | visual area 2 |
| 34.535 | 35.198 | Right | Ipsilateral | Paxinos | visual area 2 |
| 28.531 | 19.540 | Left | Contralateral | Paxinos | area 19 of cortex dorsointermediate part |
| 30.973 | 54.023 | Left | Ipsilateral | Paxinos | area 19 of cortex dorsointermediate part |
| 32.688 | 34.483 | Right | Contralateral | Paxinos | area 19 of cortex dorsointermediate part |
| 32.428 | 20.270 | Right | Ipsilateral | Paxinos | fundus of superior temporal sulcus area of cortex |
| 33.072 | 16.505 | Left | Contralateral | Paxinos | visual area 3 (ventrolateral posterior area) |
| 37.705 | 47.573 | Left | Ipsilateral | Paxinos | visual area 3 (ventrolateral posterior area) |
| 36.600 | 23.717 | Right | Contralateral | Paxinos | visual area 3 (ventrolateral posterior area) |
| 35.572 | 38.558 | Right | Ipsilateral | Paxinos | visual area 3 (ventrolateral posterior area) |
| 31.270 | 49.474 | Left | Contralateral | Paxinos | visual area 3A (dorsoanterior area) |
| 31.946 | 57.895 | Left | Ipsilateral | Paxinos | visual area 3A (dorsoanterior area) |
| 32.735 | 57.895 | Right | Ipsilateral | Paxinos | visual area 3A (dorsoanterior area) |
| 29.892 | 40.000 | Right | Contralateral | Paxinos | visual area 3A (dorsoanterior area) |
| 31.193 | 24.147 | Left | Contralateral | Paxinos | visual area 4 (ventrolatereral anterior area) |
| 36.580 | 57.743 | Left | Ipsilateral | Paxinos | visual area 4 (ventrolatereral anterior area) |
| 29.941 | 15.486 | Right | Contralateral | Paxinos | visual area 4 (ventrolatereral anterior area) |
| 33.447 | 35.958 | Right | Ipsilateral | Paxinos | visual area 4 (ventrolatereral anterior area) |
| 32.555 | 19.444 | Left | Contralateral | Paxinos | visual area 4 transitional part (middle temporal crescent) |
| 37.930 | 74.306 | Left | Ipsilateral | Paxinos | visual area 4 transitional part (middle temporal crescent) |
| 29.000 | 13.194 | Right | Contralateral | Paxinos | visual area 4 transitional part (middle temporal crescent) |
| 33.467 | 50.000 | Right | Ipsilateral | Paxinos | visual area 4 transitional part (middle temporal crescent) |
| 36.691 | 18.919 | Right | Contralateral | Paxinos | fundus of superior temporal sulcus area of cortex |
| 35.522 | 31.347 | Left | Ipsilateral | Paxinos | temporal area TE3 (inferior temporal cortex) |
| 34.795 | 13.212 | Right | Contralateral | Paxinos | temporal area TE3 (inferior temporal cortex) |
| 32.060 | 14.685 | Left | Contralateral | Paxinos | visual area 5 (middle temporal area) |
| 37.780 | 67.832 | Left | Ipsilateral | Paxinos | visual area 5 (middle temporal area) |
| 30.677 | 33.566 | Right | Contralateral | Paxinos | visual area 5 (middle temporal area) |
| 33.030 | 57.343 | Right | Ipsilateral | Paxinos | visual area 5 (middle temporal area) |
| 40.917 | 58.974 | Left | Contralateral | Paxinos | visual area 6 (dorsomedial area) |
| 40.756 | 80.057 | Left | Ipsilateral | Paxinos | visual area 6 (dorsomedial area) |
| 41.957 | 56.410 | Right | Contralateral | Paxinos | visual area 6 (dorsomedial area) |
| 35.327 | 59.829 | Right | Ipsilateral | Paxinos | visual area 6 (dorsomedial area) |

| | | | | | |
|---|---|---|---|---|---|
| 37.093 | 89.899 | Left | Contralateral | Paxinos | visual area 6A (Posterior parietal medial area) |
| 42.059 | 103.030 | Left | Ipsilateral | Paxinos | visual area 6A (Posterior parietal medial area) |
| 29.450 | 68.687 | Right | Contralateral | Paxinos | visual area 6A (Posterior parietal medial area) |
| 28.549 | 35.354 | Right | Ipsilateral | Paxinos | visual area 6A (Posterior parietal medial area) |
| 31.881 | 10.345 | Right | Ipsilateral | Paxinos | area 19 of cortex dorsointermediate part |
| 38.362 | 80.000 | Left | Contralateral | Paxinos | area 19 of cortex medial part |
| 39.798 | 109.091 | Left | Ipsilateral | Paxinos | area 19 of cortex medial part |
| 32.281 | 54.545 | Right | Contralateral | Paxinos | area 19 of cortex medial part |
| 35.660 | 77.273 | Right | Ipsilateral | Paxinos | area 19 of cortex medial part |
| 34.142 | 20.000 | Left | Ipsilateral | SAM | CA4 subfield of hippocampus |
| 30.209 | 20.000 | Right | Ipsilateral | SAM | CA4 subfield of hippocampus |
| 33.388 | 28.846 | Left | Ipsilateral | SAM | fascia dentata (granule cell layer) |
| 32.000 | 11.538 | Right | Ipsilateral | SAM | fascia dentata (granule cell layer) |
| 29.853 | 24.390 | Left | Ipsilateral | SAM | parasubiculum |
| 38.464 | 60.714 | Left | Ipsilateral | SAM | presubiculum |
| 35.070 | 10.714 | Right | Contralateral | SAM | presubiculum |
| 31.860 | 21.429 | Right | Ipsilateral | SAM | presubiculum |
| 32.968 | 61.538 | Left | Ipsilateral | SAM | prosubiculum |
| 32.771 | 15.385 | Right | Contralateral | SAM | prosubiculum |
| 36.784 | 55.556 | Left | Ipsilateral | SAM | subiculum |
| 38.336 | 27.778 | Right | Contralateral | SAM | subiculum |
| 29.215 | 13.889 | Right | Ipsilateral | SAM | subiculum |
| 30.611 | 39.655 | Left | Contralateral | Paxinos | area 13 of cortex lateral part |
| 39.979 | 32.759 | Left | Ipsilateral | Paxinos | area 13 of cortex lateral part |
| 35.960 | 27.586 | Right | Ipsilateral | Paxinos | area 13 of cortex lateral part |
| 32.184 | 43.662 | Left | Contralateral | Paxinos | area 13 of cortex medial part |
| 45.999 | 39.437 | Left | Ipsilateral | Paxinos | area 13 of cortex medial part |
| 38.520 | 14.085 | Right | Contralateral | Paxinos | area 13 of cortex medial part |
| 37.612 | 49.296 | Right | Ipsilateral | Paxinos | area 13 of cortex medial part |
| 72.378 | 39.437 | Left | Ipsilateral | Paxinos | orbital periallocortex |
| 57.053 | 53.521 | Right | Ipsilateral | Paxinos | orbital periallocortex |
| 89.477 | 65.574 | Left | Ipsilateral | Paxinos | orbital proisocortex |
| 84.582 | 70.492 | Right | Ipsilateral | Paxinos | orbital proisocortex |
| 36.348 | 39.468 | Left | Contralateral | Paxinos | primary visual cortex |
| 38.944 | 67.583 | Left | Ipsilateral | Paxinos | primary visual cortex |
| 36.093 | 52.239 | Right | Contralateral | Paxinos | primary visual cortex |
| 36.797 | 52.373 | Right | Ipsilateral | Paxinos | primary visual cortex |
| 100.000 | 112.500 | Left | Ipsilateral | Paxinos | agranular insular cortex |
| 100.000 | 87.500 | Right | Ipsilateral | Paxinos | agranular insular cortex |
| 49.854 | 18.182 | Left | Ipsilateral | Paxinos | auditory cortex anterolateral area |
| 52.670 | 18.182 | Right | Ipsilateral | Paxinos | auditory cortex anterolateral area |
| 59.362 | 13.043 | Left | Ipsilateral | Paxinos | auditory cortex caudal parabelt area |
| 29.113 | 21.739 | Right | Contralateral | Paxinos | auditory cortex caudal parabelt area |
| 46.740 | 13.043 | Right | Ipsilateral | Paxinos | auditory cortex caudal parabelt area |
| 29.467 | 13.636 | Right | Contralateral | Paxinos | auditory cortex caudolateral area |
| 44.933 | 41.667 | Left | Ipsilateral | Paxinos | auditory cortex rostral area |
| 61.304 | 44.444 | Right | Ipsilateral | Paxinos | auditory cortex rostral area |
| 62.476 | 24.390 | Left | Ipsilateral | Paxinos | auditory cortex rostral parabelt |
| 31.346 | 24.390 | Right | Contralateral | Paxinos | auditory cortex rostral parabelt |
| 80.462 | 106.667 | Left | Ipsilateral | Paxinos | auditory cortex rostromedial area |
| 88.751 | 93.333 | Right | Ipsilateral | Paxinos | auditory cortex rostromedial area |
| 65.630 | 47.826 | Left | Ipsilateral | Paxinos | auditory cortex rostrotemporal |
| 79.511 | 63.043 | Right | Ipsilateral | Paxinos | auditory cortex rostrotemporal |
| 28.448 | 15.789 | Right | Ipsilateral | Paxinos | auditory cortex rostrotemporal lateral area |
| 70.596 | 75.000 | Left | Ipsilateral | Paxinos | auditory cortex rostrotemporal medial area |
| 79.636 | 87.500 | Right | Ipsilateral | Paxinos | auditory cortex rostrotemporal medial area |
| 88.798 | 55.000 | Left | Ipsilateral | Paxinos | dysgranular insular cortex |
| 90.145 | 62.500 | Right | Ipsilateral | Paxinos | dysgranular insular cortex |
| 62.330 | 28.788 | Left | Ipsilateral | Paxinos | granular insular cortex |
| 81.489 | 30.303 | Right | Ipsilateral | Paxinos | granular insular cortex |
| 31.246 | 26.471 | Left | Contralateral | Paxinos | insular proisocortex |
| 88.721 | 64.706 | Left | Ipsilateral | Paxinos | insular proisocortex |
| 80.933 | 61.765 | Right | Ipsilateral | Paxinos | insular proisocortex |
| 84.805 | 53.571 | Left | Ipsilateral | Paxinos | parainsular cortex lateral part |
| 67.781 | 42.857 | Right | Ipsilateral | Paxinos | parainsular cortex lateral part |
| 94.439 | 86.842 | Left | Ipsilateral | Paxinos | parainsular cortex medial part |
| 94.076 | 94.737 | Right | Ipsilateral | Paxinos | parainsular cortex medial part |
| 78.396 | 66.667 | Left | Ipsilateral | Paxinos | temporal proisocortex |
| 82.665 | 77.778 | Right | Ipsilateral | Paxinos | temporal proisocortex |
| 77.928 | 46.759 | Left | Ipsilateral | Paxinos | temporo-parieto-occipital association area (superior temporal polysensory cortex) |
| 38.240 | 41.667 | Right | Contralateral | Paxinos | temporo-parieto-occipital association area (superior temporal polysensory cortex) |
| 61.920 | 54.630 | Right | Ipsilateral | Paxinos | temporo-parieto-occipital association area (superior temporal polysensory cortex) |
| 30.654 | 10.256 | Left | Ipsilateral | Paxinos | temporoparietal transitional area |
| 33.210 | 10.256 | Right | Contralateral | Paxinos | temporoparietal transitional area |
| 29.836 | 19.231 | Right | Ipsilateral | Paxinos | temporoparietal transitional area |

| | | | | | |
|---|---|---|---|---|---|
| 55.072 | 25.581 | Left | Ipsilateral | Paxinos | temporopolar proisocortex |
| 59.671 | 36.047 | Right | Ipsilateral | Paxinos | temporopolar proisocortex |
| 48.129 | 35.484 | Left | Contralateral | Paxinos | area 24a of cortex |
| 33.920 | 38.710 | Left | Ipsilateral | Paxinos | area 24a of cortex |
| 34.915 | 67.742 | Right | Contralateral | Paxinos | area 24a of cortex |
| 41.052 | 38.710 | Right | Ipsilateral | Paxinos | area 24a of cortex |
| 42.291 | 13.393 | Left | Contralateral | Paxinos | area 24b of cortex |
| 33.996 | 12.500 | Left | Ipsilateral | Paxinos | area 24b of cortex |
| 35.789 | 56.250 | Right | Contralateral | Paxinos | area 24b of cortex |
| 47.176 | 22.321 | Right | Ipsilateral | Paxinos | area 24b of cortex |
| 30.115 | 36.207 | Right | Contralateral | Paxinos | area 24c of cortex |
| 35.639 | 31.034 | Right | Ipsilateral | Paxinos | area 24c of cortex |
| 31.593 | 16.364 | Left | Contralateral | Paxinos | area 24d of cortex |
| 89.368 | 78.947 | Left | Ipsilateral | Paxinos | gustatory cortex |
| 89.758 | 65.789 | Right | Ipsilateral | Paxinos | gustatory cortex |
| 70.176 | 37.736 | Left | Ipsilateral | Paxinos | secondary somatosensory cortex parietal rostral area |
| 75.343 | 39.623 | Right | Ipsilateral | Paxinos | secondary somatosensory cortex parietal rostral area |
| 35.448 | 47.619 | Left | Ipsilateral | SAM | accessory basal nucleus of amygdala, magnocellular |
| 41.079 | 52.381 | Right | Ipsilateral | SAM | accessory basal nucleus of amygdala, magnocellular |
| 37.334 | 31.818 | Left | Ipsilateral | SAM | accessory basal nucleus of amygdala, parvicellular |
| 49.470 | 63.636 | Right | Ipsilateral | SAM | accessory basal nucleus of amygdala, parvicellular |
| 30.145 | 75.000 | Left | Ipsilateral | SAM | accessory basal nucleus of amygdala, superficial division |
| 47.106 | 75.000 | Right | Ipsilateral | SAM | accessory basal nucleus of amygdala, superficial division |
| 30.198 | 50.000 | Left | Ipsilateral | SAM | anterior amygdaloid area |
| 50.485 | 90.909 | Left | Ipsilateral | SAM | anterior cortical nucleus |
| 39.696 | 54.545 | Right | Ipsilateral | SAM | anterior cortical nucleus |
| 81.154 | 50.000 | Left | Ipsilateral | SAM | basal nucleus of amygdala, intermediate subdivision |
| 30.488 | 12.500 | Right | Contralateral | SAM | basal nucleus of amygdala, intermediate subdivision |
| 66.247 | 75.000 | Right | Ipsilateral | SAM | basal nucleus of amygdala, intermediate subdivision |
| 83.017 | 69.565 | Left | Ipsilateral | SAM | basal nucleus of amygdala, magnocellular subdivision |
| 74.331 | 73.913 | Right | Ipsilateral | SAM | basal nucleus of amygdala, magnocellular subdivision |
| 42.843 | 50.000 | Left | Ipsilateral | SAM | basal nucleus of amygdala, parvicellular subdivision |
| 46.946 | 67.857 | Right | Ipsilateral | SAM | basal nucleus of amygdala, parvicellular subdivision |
| 88.413 | 57.143 | Left | Ipsilateral | SAM | central nucleus of amygdala, lateral division |
| 80.061 | 85.714 | Right | Ipsilateral | SAM | central nucleus of amygdala, lateral division |
| 37.202 | 33.333 | Left | Ipsilateral | SAM | central nucleus of amygdala, medial division |
| 30.114 | 33.333 | Right | Ipsilateral | SAM | central nucleus of amygdala, medial division |
| 28.678 | 60.000 | Left | Ipsilateral | SAM | Cholinergic cell group 1 |
| 28.376 | 20.000 | Right | Contralateral | SAM | Cholinergic cell group 1 |
| 26.514 | 18.182 | Left | Ipsilateral | SAM | Cholinergic cell group 2 (nucleus of vertical limb of the diagonal band) |
| 27.300 | 27.273 | Right | Contralateral | SAM | Cholinergic cell group 2 (nucleus of vertical limb of the diagonal band) |
| 29.115 | 42.857 | Right | Contralateral | SAM | Cholinergic cell group 4, anterior lateral division (nucleus basalis of Meynert) |
| 51.717 | 14.286 | Left | Ipsilateral | SAM | Cholinergic cell group 4, intermedio-dorsal division (nucleus basalis of Meynert) |
| 30.826 | 10.949 | Left | Contralateral | SAM | Claustrum |
| 97.939 | 100.000 | Left | Ipsilateral | SAM | Claustrum |
| 97.511 | 93.431 | Right | Ipsilateral | SAM | Claustrum |
| 75.453 | 48.810 | Left | Ipsilateral | SAM | lateral nucleus of amygdala |
| 80.111 | 51.190 | Right | Ipsilateral | SAM | lateral nucleus of amygdala |
| 35.401 | 16.667 | Left | Ipsilateral | SAM | medial nucleus of amygdala |
| 31.397 | 66.667 | Left | Ipsilateral | SAM | nucleus of the lateral olfactory tract |
| 48.812 | 100.000 | Right | Ipsilateral | SAM | nucleus of the lateral olfactory tract |
| 47.781 | 27.273 | Left | Ipsilateral | SAM | paralaminar nucleus in amygdala |
| 33.643 | 31.818 | Right | Ipsilateral | SAM | paralaminar nucleus in amygdala |
| 43.691 | 41.667 | Right | Ipsilateral | SAM | periamygdaloid cortex 3 |
| 73.032 | 58.333 | Left | Ipsilateral | SAM | periamygdaloid cortex o/piriform cortex |
| 71.487 | 75.000 | Right | Ipsilateral | SAM | periamygdaloid cortex o/piriform cortex |
| 41.573 | 44.444 | Right | Ipsilateral | SAM | periamygdaloid cortex, sulcal portion |
| 32.025 | 25.000 | Left | Ipsilateral | SAM | posterior cortical nucleus |
| 40.558 | 47.305 | Left | Ipsilateral | Paxinos | medial superior temporal area of cortex |
| 34.670 | 55.689 | Right | Contralateral | Paxinos | medial superior temporal area of cortex |
| 35.215 | 34.731 | Right | Ipsilateral | Paxinos | medial superior temporal area of cortex |
| 58.105 | 11.111 | Left | Ipsilateral | Paxinos | secondary somatosensory cortex parietal ventral area |
| 50.077 | 24.074 | Right | Ipsilateral | Paxinos | secondary somatosensory cortex parietal ventral area |
| 94.066 | 26.613 | Left | Ipsilateral | SAM | anterior commissure |
| 33.106 | 13.710 | Right | Contralateral | SAM | anterior commissure |
| 90.349 | 30.645 | Right | Ipsilateral | SAM | anterior commissure |
| 27.049 | 14.286 | Left | Ipsilateral | SAM | brachium of inferior colliculus |
| 29.126 | 19.231 | Right | Contralateral | SAM | commissure of superior colliculus |
| 46.634 | 10.694 | Right | Ipsilateral | SAM | internal capsule, cerebral peduncle, corticospinal, or corticobulbar tract |
| 31.657 | 14.286 | Right | Contralateral | SAM | mammillothalamic tract |
| 28.855 | 11.650 | Right | Ipsilateral | SAM | middle cerebellar peduncle |
| 28.518 | 10.417 | Right | Contralateral | SAM | Muratoff bundle |
| 33.942 | 10.417 | Right | Ipsilateral | SAM | Muratoff bundle |
| 26.762 | 14.286 | Right | Contralateral | SAM | stria medullaris |
| 30.689 | 18.182 | Right | Ipsilateral | SAM | superior cerebellar peduncle decussation |

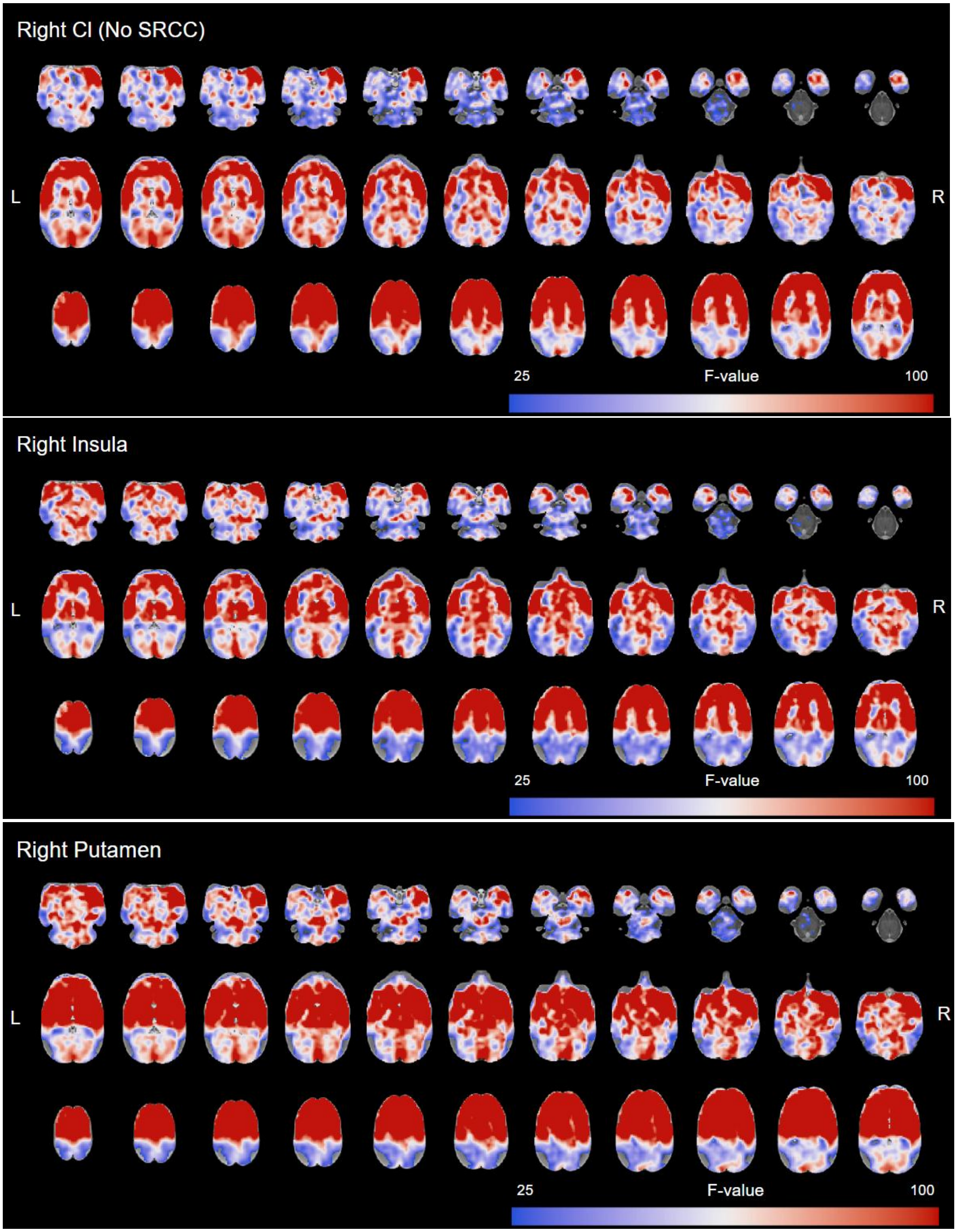
Right Cl (No SRCC)
L
R
25
F-value
100
Right Insula
L
R
25
F-value
100
Right Putamen
L
R
25
F-value
100

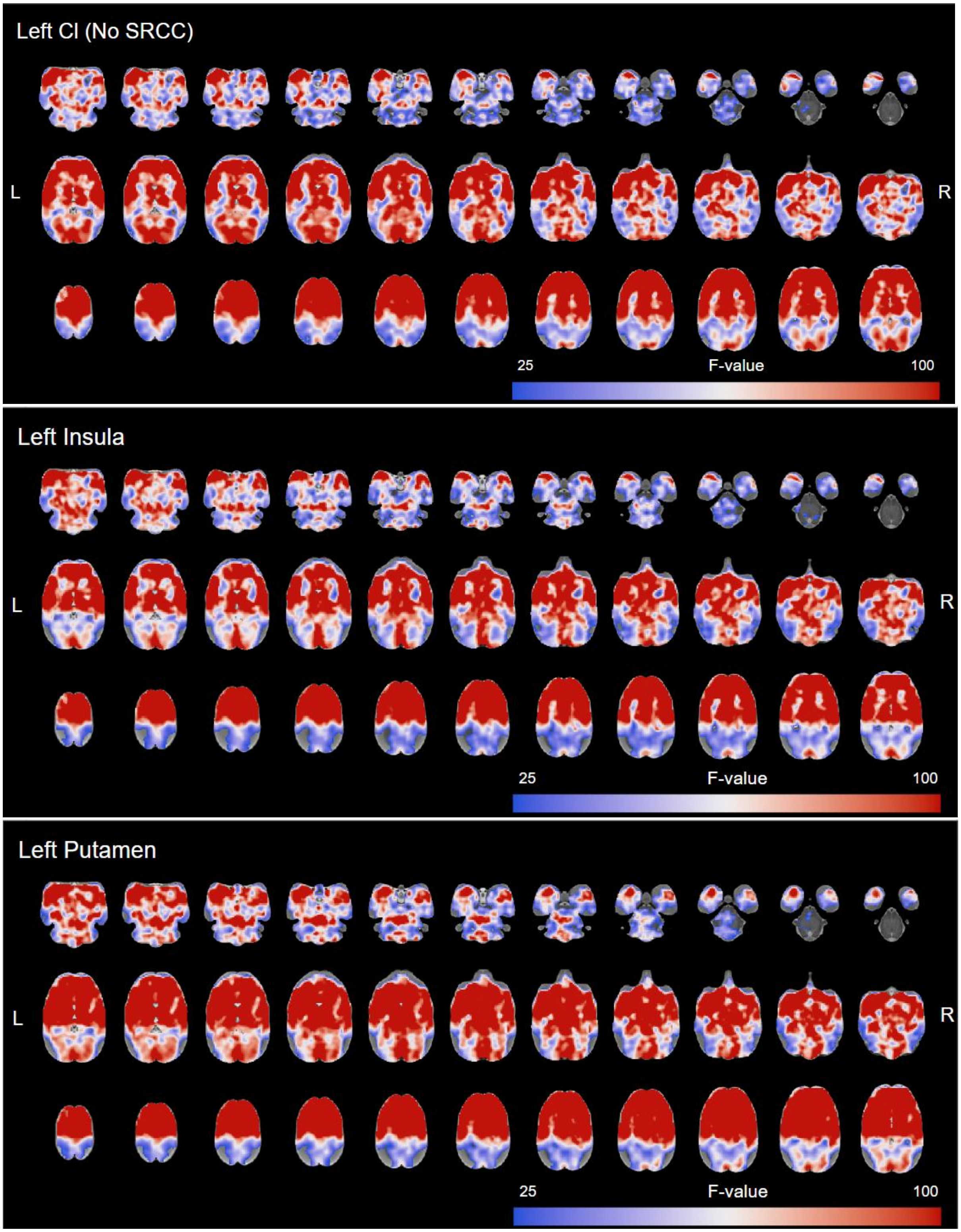


**Figure 1.** Resting state functional connectivity maps using the right and left claustrum pre-SRCC, insula, putamen as seeds. Data presented (n=28) as significant F values (>25.987)
